# Redox-Triggered Coupling Network Mediates Long-Range Energy Transduction in Respiratory Complex I

**DOI:** 10.64898/2026.04.14.716442

**Authors:** Niklas L. Hoja, Jonas Hentschel, Hyunho Kim, Thilo Seifermann, Adel Beghiah, Tim Schlosser, Patricia Saura, Thorsten Friedrich, Ville R. I. Kaila

**Affiliations:** Department of Biochemistry and Biophysics, Stockholm University, 10691, Stockholm, Sweden; Institut für Biochemie, Albert-Ludwigs-Universität Freiburg, Germany

**Keywords:** cellular respiration, energy conversion, proteoliposomes, proton transfer, QM/MM, cryo-EM

## Abstract

Complex I is a gigantic redox-driven proton pump that powers oxidative phosphorylation by a unique long-range (>200 Å) proton-coupled electron transfer process. To elucidate the molecular principles underlying this intricate *action-at-a-distance* effect, we combine here multiscale quantum/classical (QM/MM) simulations with site-directed mutagenesis, proteoliposome experiments, and cryo-electron microscopy (cryo-EM). We find that quinol binding at a distinct site within a local membrane cavity, triggers a long-range protonation cascade over water-mediated proton wires along a conserved carboxylate pathway (E-channel). We identify a central mechanical switch point, comprising the conserved Tyr156^H^, the mutation of which impedes conformational changes along conserved loops, but not the proton transfer reaction itself. Using our integrative multi-disciplinary approach, we reveal central coupling sites along the redox-driven proton transport process mediating energy conversion in Complex I, and illustrate the power of theory-guided experiments.

## Introduction

Respiratory Complex I (NADH:ubiquinone oxidoreductase) is the largest (0.5-1 MDa) and most intricate enzyme of aerobic electron transport chains^1–4^. Complex I catalyses electron transfer from nicotinamide adenine dinucleotide (NADH) to ubiquinone (Q), and couples this to translocation of protons across the inner mitochondrial or bacterial cytoplasmic membrane (Fig. 1). Its proton pumping activity generates a proton motive force (PMF) that powers the synthesis of adenosine triphosphate (ATP)^5,6^, whilst the reduced quinol pool shuttles the electron *via* Complex III (cytochrome *bc*_1_) to Complex IV (cytochrome *c* oxidase), reducing dioxygen (O_2_) to water (H_2_O)^7,8^. Complex I is conserved across bacteria and mitochondria, while its dysfunction is implicated in numerous human mitochondrial disorders^9^.

**Fig. 1.**
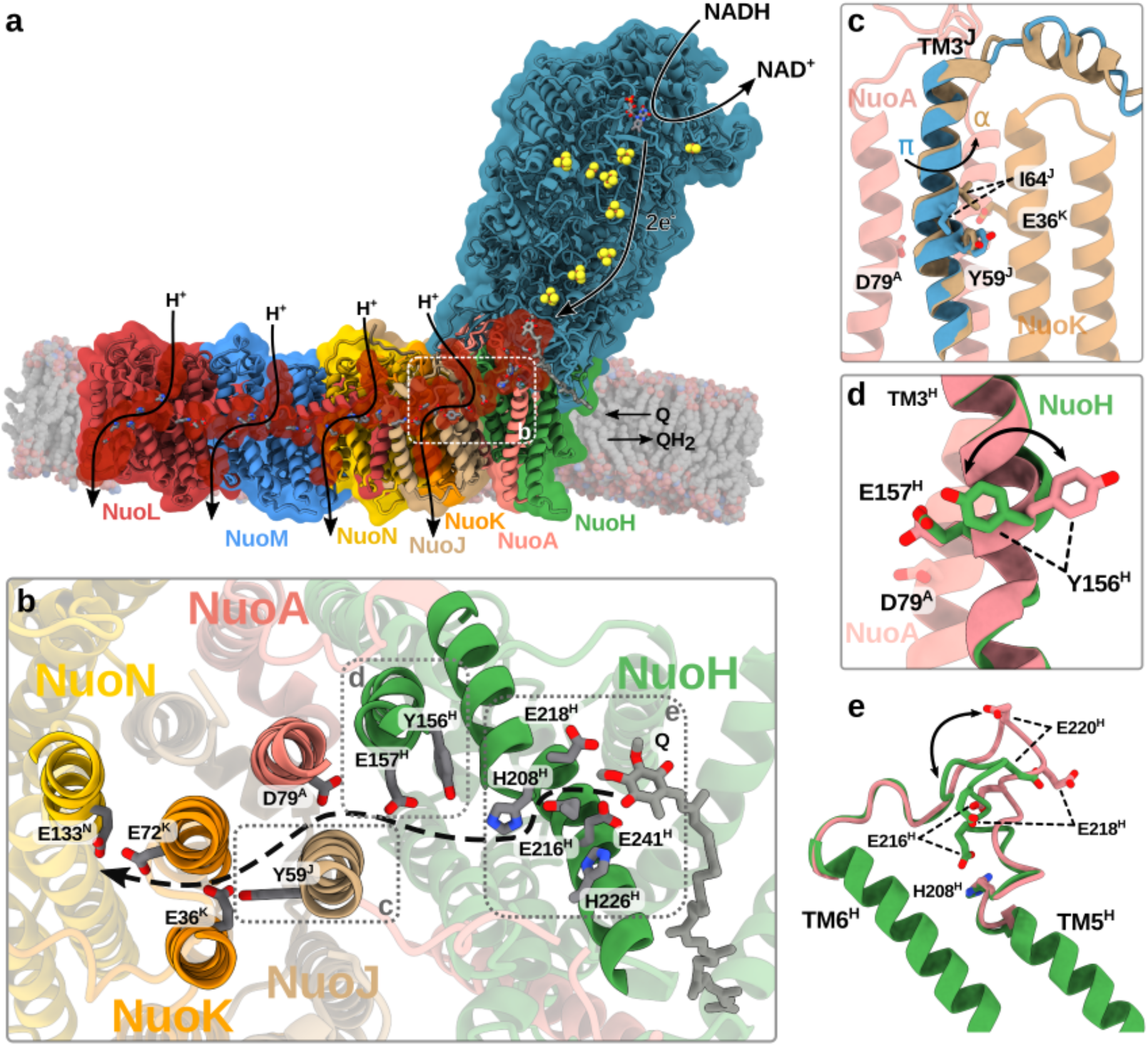
Structure and function of Complex I. **a**, NADH oxidation drives long-range electron transfer through the hydrophilic domain, which results in reduction of quinone (Q) to quinol (QH_2_). The redox activity triggers proton pumping across the 200 Å long membrane domain (PDB ID: 7Z7S^12^). **b**, Closeup of the E-channel leading from Q site 2 *via* a cluster of carboxylates to the interface of NuoN (top view). **c-e**, Functional elements linked to the activation of the E-channel and the redox-driven proton transport process: **c**, π→α switching of TM3^J^; **d**, *outward (pink)* → *inward (green)* flip of Tyr156^H^; **e**, conformational change of the TM5-6 loop of NuoH.

The hydrophilic domain of Complex I mediates NADH-driven electron transfer *via* flavin mononucleotide (FMN) and a chain of iron-sulphur (FeS) clusters to Q, which binds at the end of a *ca*. 40 Å long substrate tunnel at the interface of the hydrophilic and membrane domains (Fig. 1). The reduction of Q to ubiquinol (QH_2_) results in proton pumping across a remarkable distance of >200 Å within the membrane domain in a process that is highly efficient and fully reversible. Yet, despite major structural, biochemical, and computational efforts^10–19^, the precise long-range coupling mechanism remains elusive, and one of the most central open questions in bioenergetics.

Proton pumping in Complex I is catalysed by the four putative proton translocation pathways in a bundle established by the NuoH/A/J/K subunits and in the antiporter-like subunits NuoN, NuoM, and NuoL (*E. coli* nomenclature) that form a functionally critical proton-conducting network. The proton pumping is believed to be initiated by redox-triggered protonation and conformational changes within the E-channel, a carboxylate-rich region located around the Q tunnel, and extending along the membrane domain *via* the NuoH/A/J/K region towards NuoN (Fig. 1a,b). Recent studies^11,12,20,21^ revealed conformational changes around conserved loops and in trans-membrane helix 3 of NuoJ (TM3^J^), which transitions from a π-helix to an α-helical form upon activation of Complex I (Fig. 1c,d)^22,23^, modulating the hydration state of the pathway^11,20^ and lowering the barrier for directional proton transfer^21^. A similar α→π transition, together with conformational changes in surrounding loops were also linked to the *deactive* (D) form of mammalian isoforms^24^, which may represent an off-cycle regulatory state^13,25^ that protects against formation of reactive oxygen species (ROS) upon reverse electron transfer (RET)^26,27^. Accordingly, the E-channel and the surrounding flexible loops could serve as a conformational hub that connects the local redox chemistry with the global propagation of a charge wave across the membrane domain, as suggested in the electrical wave mechanism (*cf*. Refs.^1,21^). Despite recent progress in understanding the elementary steps of this process, the molecular principles remain much debated and controversial^12,16,17,28,29^.

The E-channel forms a water-filled cavity, starting from the nearby membrane-bound Q binding site (Q site 2, Fig. 1b), where the quinol is proposed to bind after its reduction and protonation in the active site (Q site 1), near the N2 FeS centre^30–32^. The E-channel continues *via* Glu216^H^/Glu218^H^/Glu220^H^ and Asp213^H^ to His208^H^, and further along Tyr156^H^ and Glu157^H^ to Asp79^A^ (*E. coli* numbering) that support Grotthuss-type water-mediated proton transfer reactions^12,21^. Asp79^A^ was recently suggested to function as a central kinetic gate^33^, by establishing a water-mediated array with Glu36^K^ and Glu72^K^, at the interface to NuoN, in the α-state of TM3^J^ (Fig. 1b,c), whilst the π-conformation of TM3^J^ blocks the hydration and proton transfer by steric clashes^11,12,20,21^. It was also proposed that the E-channel could serve as a direct source of protons in the Q reduction process^12^, although there is currently no direct experimental evidence supporting this proposal. Nevertheless, mutation of residues in the E-channel perturbs or entirely abolishes the coupled proton-electron transfer activity^33–37^, thus indirectly supporting its central role in mediating the long-range coupling effects.

The π/α-transition of TM3^J^ and activation of the E-channel has also been linked to the conformational state of the conserved Tyr156^H^ of TM4^H^ (Fig. 1d)^12,20,21^. In the α-helical form of TM3^J^ (*viz*. the *‘active’* state), Tyr156^H^ is orientated towards the hydrated E-channel (*inward* conformation). In contrast, in the ‘*resting*’^12^ and ‘*deactive’* forms of Complex I^11,22^, with TM3^J^ in the π-bulge form, Tyr156^H^ rotates away from the E-channel (Fig. 1b,d). These conformational changes are conserved across Complex I from various species, thus suggesting that the conformational switching of Tyr156^H^ could have a functional role in the long-range charge propagation, *e*.*g*., by controlling the proton transfer activity along the E-channel, as suggested by Sazanov and co-workers^12^. Nevertheless, the molecular principles underlying this process remain much debated, and direct experimental evidence supporting its involvement in the proton transfer reactions is lacking. To systematically test the possible role of Tyr156^H^, we combine here multiscale quantum and classical simulations with free energy explorations to guide site-directed mutagenesis experiments, proton pumping assays in proteoliposomes, and cryoelectron microscopy (cryo-EM).

## Results

### Mechanism of water-mediated proton transfer along the E-channel

To explore the molecular principles underlying the Tyr156^H^ flip, we first performed atomistic molecular dynamics (MD) simulations of the *E. coli* Complex I, based on the active state cryo-EM structure (PDB ID: 7Z7S^12^) with an α-helical TM3^J^, and a quinol or QH^-^ species (Q_8_) modelled in Q site 2, based on previous simulations of Complex I^20,21^ (see *Methods*, Supplementary Fig. 1). We embedded the model into a three-component (POPE/POPG/cardiolipin) lipid membrane, ions, and water molecules, resulting in a system with around 0.9 million atoms. The system was propagated for a cumulative simulation time of *ca*. 10 μs (with 0.5 μs trajectories of individual states, Supplementary Table 1). Our MD simulations suggest that the quinol binds in Q site 2 by a π-cation interaction with Arg87^B^ together with hydrogen-bonds with a surrounding cluster of conserved carboxylates on TM5-6 loop of NuoH (TM5-6^H^) (Fig. 2a, Supplementary Fig. 3). In particular, Glu216^H^, which was previously implied as a transient acceptor of a quinol proton (*cf*. Refs.^20,21^), stabilises the quinol by hydrogen-bonded interaction. However, we note that other conserved carboxylates near the entrance of the E-channel and the quinol (*e*.*g*. Glu218^H^) could serve a similar function, indicating possible functional cooperativity and/or redundancy. Upon proton transfer from the quinol to Glu216^H^ (*cf*. Ref.^21^), forming a QH^-^/GluH state, the protonated carboxylate rapidly rotates away from Q site 2 and forms a water-mediated contact with His208^H^ (Fig. 2a, Supplementary Figs. 3 and 4). In turn, His208^H^ hydrogen-bonds *via* a water molecule to Tyr156^H^ and Glu157^H^ (Fig. 2b, Supplementary Fig. 4), although sub-populations of other water-mediated contacts bypassing Tyr156^H^ and/or His208^H^ are observed as well (Supplementary Fig. 4). The *inward* conformation of Tyr156^H^ together with Thr153^H^ (Fig. 2b, Supplementary Figs. 2 and 4), stabilises the water wire *via* hydrogen-bond contacts.

**Fig. 2.**
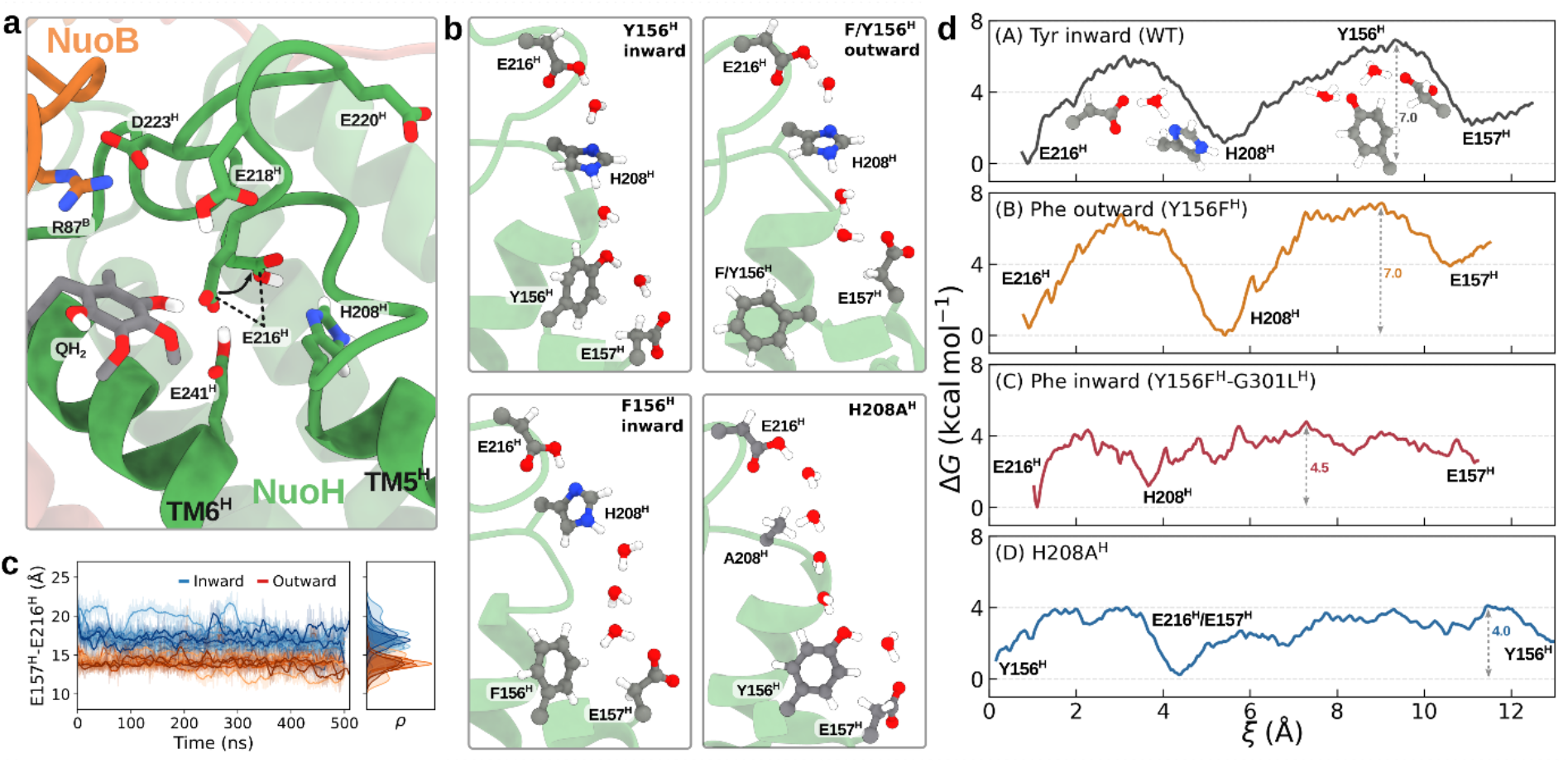
Multiscale simulations of long-range proton transfer along the E-channel. **a**, Quinol binding to Q site 2 triggers proton transfer to Glu216^H^, and formation of a proton conduit along the E-channel. **b**, Water chain formed in different conformational states of Tyr156^H^/Phe156^H^ from MD simulations. **c**, Distance between Glu216^H^ and Glu157^H^ from MD simulations, with *inward* (in *blue*) and *outward* (in *orange*) Tyr156^H^. **d**, Free energy profile of proton transfer with (A) *inward* Tyr156^H^, (B) *outward* Phe156^H^, (C) *inward* Phe156^H^, and (D) the H208A^H^ variant.

To probe the energetics of the proton transfer reaction along the pathway formed between Glu216^H^ and Glu157^H^, we next applied hybrid quantum/classical (QM/MM) free energy simulations (see *Methods*, Supplementary Figs. 1b and 5), to accurately describe the energetics of the reaction at the density functional theory (DFT) level, with electronic polarization effects from the surrounding protein environment treated by a classical electrostatic embedding scheme. Our QM/MM free energy simulations suggest that the proton transfer along the Glu216^H^-His208^H^-Glu157^H^ array takes place *via* a Grotthuss-type proton hopping process, involving transient Zundel (H_5_O_2_^+^), Eigen (H_3_O^+^), and imidazolium (HisH^+^) intermediates, with Tyr156^H^ mediating a concerted proton exchange between His208^H^ and Glu157^H^ (Fig. 2b,d, Supplementary Fig. 5a, Supplementary Movie 1). The free energy barrier for this proton transfer process is *ca*. 7 kcal mol^-1^, comparable to a nanosecond transition (based on transition state theory), whereas the overall reaction is endergonic by a few kcal mol^-1^ (Fig. 2d), indicating that the reaction is both kinetically and thermodynamically feasible. A directed electric field forms from Glu216^H^ to Glu157^H^, suggesting that the proton transfer reaction is driven by electrostatic effects (Supplementary Fig. 6), as observed in other bioenergetic systems^38–43^.

To examine the functional role of the phenolic group in the proton transfer, we next constructed the Y156F^H^ variant by *in silico* mutation. Atomistic MD simulations suggest that Phe156^H^ readily adopts the *outward*-facing conformation (Supplementary Fig. 7a-c), as also supported by our free energy simulations of the conformational flip (Fig. 3c). In the *outward* state, His208^H^ hydrogen-bonds with Glu157^H^ *via* 2-3 water molecules (Fig. 2b, Supplementary Fig. 4d), enabled by a subtle conformational change of TM4^H^ that shifts Glu157^H^ towards Glu216^H^/His208^H^ (Fig. 2b). Interestingly, removal of the phenolic group along the pathway has only a negligible effect on the free energy barrier for proton transfer relative to the parent enzyme (WT, Fig. 2d, Supplementary Fig. 5b). Taken together, our multiscale simulations suggests that while the *inward* conformation of Tyr156^H^ energetically supports proton transfer, the reaction does not critically depend on this residue and remains both kinetically and thermodynamically feasible even in its absence.

**Fig. 3.**
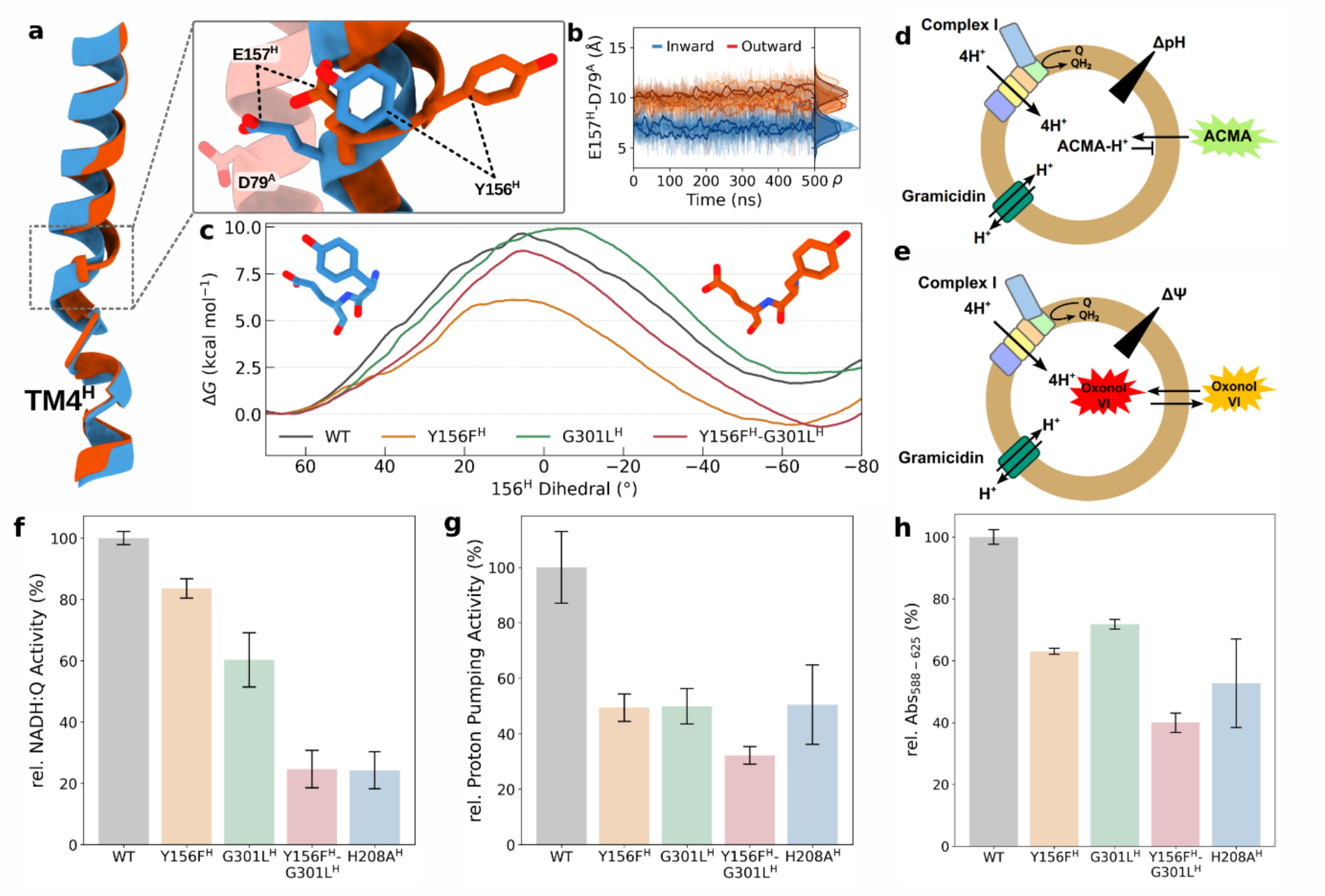
Probing the redox-coupled proton pumping activity. **a**, Structural overlay of the active (*blue*) and resting (*orange*) conformations of Complex I, highlighting the structural differences of TM4^H^, Tyr156^H^, and Glu157^H^. **b**, Distance between Glu157^H^ and Asp79^A^ from unbiased MD simulations clustered into *inward* (*blue*) and *outward* (*orange*) conformations of Tyr156^H^. **c**, Classical free energy sampling of the Tyr156^H^ flip in parent Complex I (WT) and variants. **d-e**, Scheme of the experimental setup to determine **d**, ACMA fluorescence and **e**, oxonol-VI absorbance in proteoliposomes. **f**, NADH:DQ oxidoreductase activity (100% activity of WT corresponds to 2.69 ± 0.05 U mg^-1^) and **g, h**, proton pumping activity of parent Complex I (WT) and the Y156F^H^, G310L^H^, Y156F^H^/G310L^H^, and H208A^H^ variants reconstituted into proteoliposomes. Formation of the ΔpH is probed by ACMA fluorescence, and the membrane potential is probed by oxonol VI absorbance.

To further explore the conformational switching mechanism, we performed MD simulations with Tyr156^H^ in the *outward* state (Supplementary Fig. 7a, see *Methods*), in which the residue points towards the hydrophobic cavity between TM3^H^ and TM8^H^. Notably, as in the Y156F^H^ variant, the *outward* Tyr156^H^ induces a subtle rotation of the TM4^H^, triggering a conformational change in Glu157^H^ that increases its distance from Asp79^A^ (Fig. 3a,c). Conversely, in the *inward* Tyr156^H^, the two carboxylates move closer together (by *ca*. 3 Å, Fig. 3a,b), a configuration that could favour the subsequent proton transfer from Asp79^A^ towards the interface of NuoN, as suggested by our previous work^21,33^. Interestingly, comparison of cryo-EM structures in the α- and π-states of TM3^J^ in different organisms reveals a similar repositioning of Glu157^H^ that correlates with the *inward*/*outward* flip of Tyr156^H^, thus supporting that our computational predictions are robust (Supplementary Fig. 8).

With aims to sterically hinder the conformational switching of Tyr156^H^, we next substituted Gly301^H^ on TM8^H^ into a bulky hydrophobic residue (G301L^H^) to partially block the cavity. Our classical free energy simulations of the G301L^H^ variant suggest that the barrier for the *inward → outward* flip is largely unchanged as compared to parent Complex I (WT, Δ*G*^‡^ ≈ 10 kcal mol^-1^), while the *outward* conformation is slightly disfavoured (Δ*G* ≈ 3 kcal mol^-1^, Fig. 3c). Based on these observations, we also investigated the Y156F^H^/G301L^H^ variant, which shows a similar stabilising effect of the *outward* Phe156^H^, but associated with a significantly higher barrier (Δ*G*^‡^_Y156F/G301L_ ≈ 9 kcal mol^-1^) and a slightly shifted *outward* position of Phe156^H^ (by *ca*. -10º degrees, Fig. 3c). The *inward* conformation in the Y156F^H^ variants result in an increased length of the water wire connecting His208^H^ with Glu157^H^ (Supplementary Fig. 4g,h). Surprisingly, QM/MM simulation of the Y156F^H^/G301L^H^ variant (with the *inward* Phe156^H^) shows a modest proton transfer barrier of *ca*. 5 kcal mol^-1^, thus lower than that of the WT Complex I (Fig. 2d, Supplementary Fig. 5). This suggests that, although the Y156F^H^/G301L^H^ substitution destabilises the water connectivity between His208^H^ and Glu157^H^ (Fig. 2b, Supplementary Fig. 4g,h), this variant also kinetically supports proton transfer along the E-channel.

To assess the potential role of His208^H^ in the proton transfer reaction, we also created the H208A^H^ variant *in silico*. MD simulations suggest that this variant forms a longer water chain with four water molecules, directly connecting Glu216^H^ with Tyr156^H^ (instead of three in WT Complex I), which in turn is hydrogen-bonded *via* one water molecule to Glu157^H^ (Fig. 2b, Supplementary Fig. 4i). Interestingly, the proton transfer barrier of this variant is drastically lower as compared to that of the WT (Δ*G*^‡^<4 kcal mol^-1^), with the transition state formed by a Zundel-like species between Glu216^H^ and Tyr156^H^ (Supplementary Fig. 5). In addition to the lowered proton transfer barrier, we note that the absence of His208^H^ could destabilize the *inward* position of Tyr156^H^, by removing its surrounding hydrogen-bonding network (Fig. 2b). However, despite its potential role in the *E. coli* Complex I, we note that His208^H^ is not fully conserved in all species (Supplementary Fig. 8).

Taken together, our multiscale simulations suggest that proton transfer along the E-channel from Q site 2 to Glu157^H^ is catalysed by a water-mediated proton transfer reaction with a low free energy barrier of Δ*G*^‡^<7 kcal mol^-1^, comparable to ns-μs reaction rate. Moreover, our *in silico* mutagenesis suggests that proton transfer can also take place in the absence of Tyr156^H^, or with the phenolic group in the *outward* conformation. This indicates that the Tyr156^H^ flip could, in contrast to previous proposals^12^, have an alternative role in the catalytic mechanism of Complex I, by promoting local rearrangements within the surrounding residue network (see below).

### Redox activity and proton pumping are only modestly blocked by mutation of the Tyr-switch

To experimentally probe the effects of the *in silico* mutations, we introduced the corresponding substitutions into the *E. coli* Complex I, and assessed the coupled NADH:decylquinone (DQ) oxidoreductase and proton pumping activities in proteoliposomes with ΔpH and Δψ sensitive dyes 9-amino-6-chloro-2-methoxyacridine (ACMA) and oxonol VI, respectively (Fig. 3, Supplementary Figs. 10 and 11, see *Methods*). All variants were fully assembled containing all subunits (Supplementary Fig. 10).

The NADH:DQ activity of the Y156F^H^ variant corresponds to 83 ± 3 % activity relative to the parent enzyme (WT, Fig. 3f), whereas the proton translocation is reduced to 49 ± 5 % in the ACMA assay (monitoring ΔpH) and 63 ± 1 % in the oxonol assay (monitoring Δψ), respectively (Fig. 3g,h). This subtle decrease in activity thus supports our QM/MM data, demonstrating that Tyr156^H^ is not essential for proton translocation. To further probe the underlying effects, we also studied the electron transfer and proton pumping activities of the G301L^H^ and Y156F^H^/G301L^H^ variants, which show respective NADH:DQ activities of 60 ± 8

% and 24 ± 6 %, while reducing the proton translocation activities to 50 ± 6 % (G301L^H^) and 32 ± 3 % (Y156F^H^/G301L^H^) (Fig. 3f,g). Moreover, our proton pumping assays monitored with oxonol VI suggest that all variants establish and maintain an electrochemical gradient across the membrane (ΔΨ) (Fig. 3h). As the G301L^H^ mutation does not alter any titratable residue, the lowered activity of this variant is likely to arise from changes in the local structure and/or dynamics around Tyr156^H^. In contrast, the H208A^H^ variant, which resulted in a drastic loss of proton transfer barriers in our QM/MM simulations (Fig. 2d), shows a rather strong effect in the NADH:DQ (24 ± 6%) and proton pumping (51 ± 14%) activities (see *Discussion*).

Taken together, our combined data show that the redox-coupled proton pumping activity of Complex I is only modestly diminished by the Y156F^H^ mutation or substitutions in its immediate surroundings. Mechanistically, these findings show that the *inward* → *outward* transition of Tyr156^H^ is not strictly required for the proton transfer between Glu216^H^ and Glu157^H^. Moreover, the G301L^H^ mutation has a stronger impact on both activities than the absence of a phenolic group alone, possibly as the substitution interferes with the conformational switching dynamics of Tyr156^H^. Our findings therefore point to an alternative function of Tyr156^H^ as a conformational switch point rather than a direct element responsible for proton transfer.

### Cryo-EM structures of the Tyr-variants highlight a conformational switching network

To obtain structural insights into the modifications at or around Tyr156^H^, we next determined the cryo-EM structures of the variants to local resolutions of 2.2–2.9 Å (Fig. 4a-c, Supplementary Figs. 12-16, Supplementary Table 4). All structures confirm the presence of the corresponding substitutions, with clear sidechain densities observed at the respective positions (Fig. 4b,c). The overall structure of the variants resembles previously described resting-state structures of *E. coli* Complex I^12,33^, resolved with TM3^J^ in the π-bulge conformation, but revealing conformational changes around the mutation sites and functional loops surrounding the Q cavity (Fig. 4a-c, Supplementary Fig. 16).

**Fig. 4.**
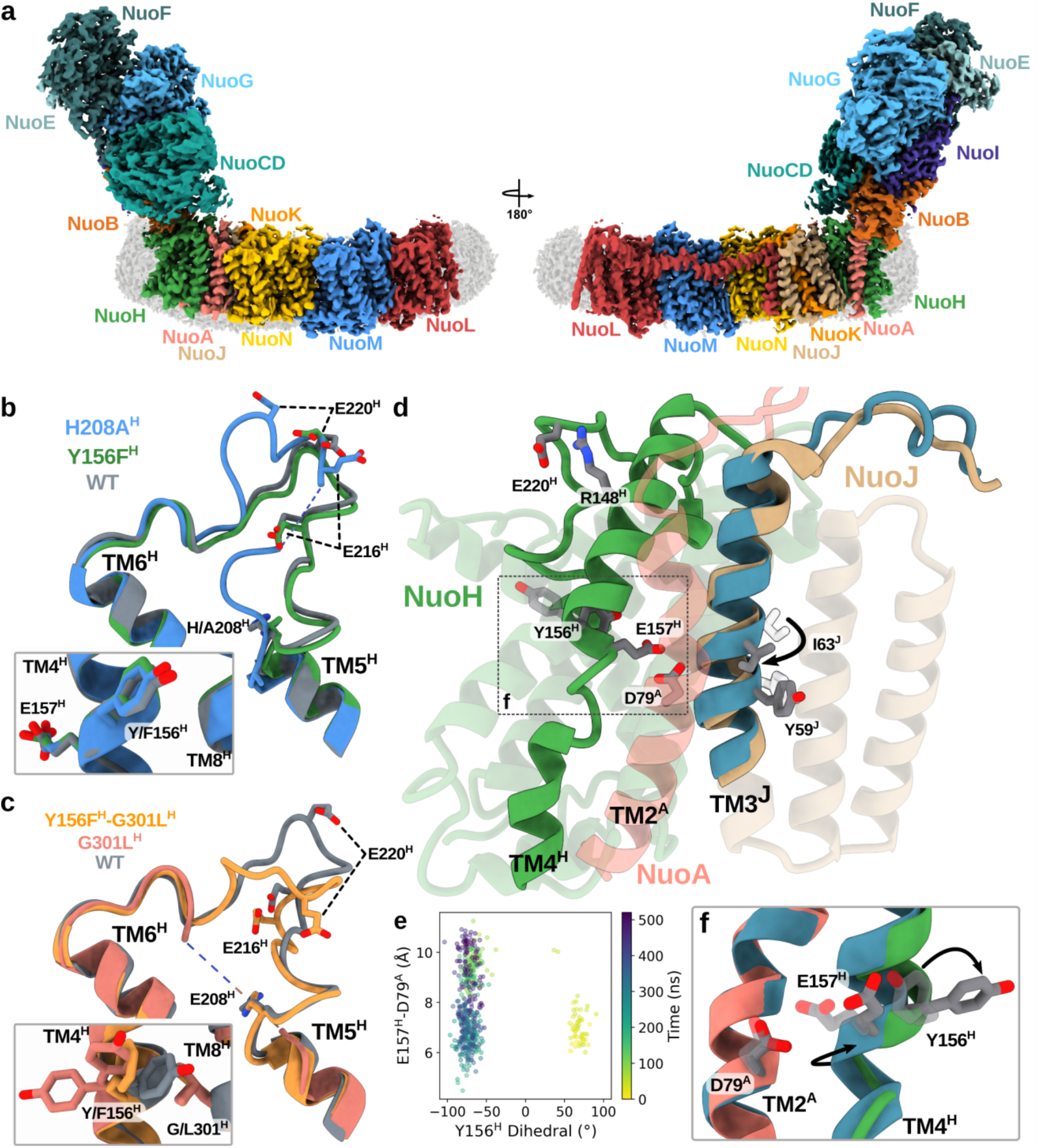
Structural exploration of the conformational switching network. | **a**, Cryo-EM map of the Y156F^H^/G301L^H^ variant shown in two side views rotated by 180°, with individual subunits labelled (see Supplementary Figs. 11-14 for other variants). **b**, Structure of the TM5-6^H^ loop in parent Complex I (WT, PDB: 9TAK^33^), Y156F^H^, and H208A^H^ variants. *Inset:* Closeup of Tyr156^H^ and Glu157^H^ in the *outward* state. **c**, Structure of the TM5-6^H^ loop in parent Complex I (WT), G301L^H^, and Y156F^H^/G301L^H^ variants *Inset:* Closeup of Tyr156^H^ and Glu157^H^, shows mixed *inward* and *outward* states, as well as a shifted *outward* position. **d**, Superimposed structures from TMD simulations, highlighting structural differences in the α-(in *beige*) and π conformations (in *blue*) of TM3^J^. **e**, The Glu157^H^-Asp79^A^ distance plotted against the Tyr156^H^ dihedral (N-Cα-Cβ-Cγ) for the duration of the TMD and the subsequent unbiased MD simulation (total: 500 ns). **f**, Repositioning of Glu157^H^ during the *inward* → *outward* flip of Tyr156^H^.

In the Y156F^H^ variant (PDB ID: 28WM), Phe156^H^ shows an *outward* conformation (Fig. 4b), whereas the TM5-6^H^ loop adopts a conformation similar to other resting state structures (Fig. 4b, but see below). Interestingly, the G301L^H^ variant (PDB ID: 28VI) shows densities of both the *inward* and *outward* conformation of Tyr156^H^ (Fig. 4c), while TM3^J^ adopts a π-bulge conformation (Supplementary Fig. 16). In contrast to the Y156F^H^ variant, the TM5-6^H^ loop is poorly resolved in our G301L^H^ structure (Fig. 4c, Supplementary Fig. 16), with missing density for the 205^H^ – 225^H^ region. The G301L^H^ substitution thus introduces a pronounced steric hindrance, shifting the *outward* position of Tyr156^H^, and creating disorder in the nearby loops (*e*.*g*. TM5-6^H^ and TM1-2^A^). Consistent with our MD simulations, the mixed *inward*/*outward* populations lead to a subtle shift of TM4^H^ and Glu157^H^ (Supplementary Fig. 17).

Remarkably, in contrast to our other variants, the TM5-6^H^ loop in Y156F^H^/G301L^H^ (PDB ID: 28TL) adopts distinct conformation, possibly an intermediate between the *active* and *resting* forms, despite the *outward* orientation of Phe156^H^ and the π-bulge at TM3^J^ (Fig. 4c, Supplementary Fig. 16). This structural difference suggests a perturbed coupling within the loops around the Q cavity and TM3^J^, which in turn, affects the conformational switching of Phe156^H^ (Fig. 4f). In this regard, it is possible that Phe156^H^ exerts less structural strain on the TM5-6^H^ loop, allowing it to locally adopt a distinct conformation even in the presence of the π-bulge at TM3^J^, while the absent phenolic group destabilizes the *inward* conformation. Supporting these findings, the H208A^H^ variant (PDB ID: 28NI), determined at 2.3 Å resolution, shows a loss of density of the TM5–6^H^ loop indicating, dynamic perturbation of this region. This destabilization is likely caused by the absence of the polar interactions normally provided by His208^H^ in parent Complex I (WT).

Our combined observations support that the Tyr156^H^ flip forms a critical element coupled to a broader conformational switching network. Indeed, analysis of B-factors from our cryo-EM data together with the *root-mean-square fluctuations* (RMSF) from our MD simulations, qualitatively support an increased loop mobility in this region (Supplementary Fig. 18).

In summary, the presence of both *inward* and *outward* orientations of Tyr156^H^ in the G301L^H^ variant, together with the unresolved TM5-6^H^ loop, suggest that the conformational state of Tyr156^H^ is strongly influenced by the local structural arrangement in conserved loops and helices. Supporting this hypothesis, the Y156F^H^/G301L^H^ variant displayed an *active*-like conformation at the TM5-6^H^ loop, despite the *outward* conformation of Phe156^H^. Our data also suggested that the conformational switching is linked to a structural re-positioning of Glu157^H^, which could favour proton transfer from Asp79^A^ to Glu36^K 33^. Our combined findings thus support a key role of Tyr156^H^ in establishing a conformational switching network that mediate redox-driven structural changes from the active site, and transducing free energy for proton translocation in the membrane domain (see *Discussion*).

### The Tyr156^H^ flip couples to the α/π-transition of TM3^J^

To explore the coupling network involving the conformational switch of Tyr156^H^, we next performed targeted MD (TMD) simulations by driving the α→π transition of TM3^J^ helix using the cryo-EM structure of Complex I in the π state as a template (PDB: 7ZCI^12^, see *Methods*). During the simulations, TM3^J^ switches into the π-conformation, and induces a spontaneous *outward* flip of Tyr156^H^, thus supporting our hypothesis that the Tyr switch couples to the α → π transition (Fig. 4d-f, Supplementary Fig. 9, Supplementary Movie 2). Remarkably, the *inward* → *outward* flip of Tyr156^H^ also results in a conformational shift of Glu157^H^, increasing its distance to Asp79^A^ (Fig. 4e,f), consistent with our unbiased MD (Fig. 3b) and cryo-EM data (Supplementary Fig. 8, 17). The α→π transition also couples to conformational changes in the TM5−6^H^ loop and repositioning of Glu220^H^ (Supplementary Fig. 9), although this region is located *ca*. 35 Å away at the interface between the hydrophilic and membrane domains. During this process, Gln214^H^ (TM5-6^H^) and Gln152^H^ (TM3^H^) undergo coupled conformational changes, which are linked to the coordinated re-arrangement of Tyr156^H^ (Supplementary Fig. 9). We note that the Glu220^H^-Arg148^H^ interaction could have a critical role for the oxidoreductase activity, *via* its connection to the β1-β2 loop of NuoCD, the Q site 2 through Arg407^CD^, and the proton transfer-mediating residues within the E-channel (Fig. 4d, Supplementary Fig. 9). Taken together, our TMD simulations support a role of Tyr156^H^ as a ‘mechanical latch’ in establishing the long-range coupling network in Complex I.

## Discussion

We have addressed here the functional role of the conformational flip of Tyr156^H^ in redox-driven proton pumping of Complex I by combining computational, biochemical, and structural experiments. We show that Tyr156^H^ is not essential for the proton transfer itself, with the conservative tyrosine substitution by a phenylalanine resulting in only a moderate (∼20-40%) reduction of the redox-coupled pumping activity. We therefore suggest that the Tyr156^H^ flip could be mediated by redox-driven processes in the Q site, linked to the π → α transition of TM3^J^.

Our TMD simulations (Fig. 4d-f, Supplementary Fig. 9) in combination with structural studies of *in silico*-designed mutations (Supplementary Fig. 7, Supplementary Table 1), suggest that the *outward* → *inward* flip of Tyr156^H^ is mediated by conformational coupling *via* conserved loops at the interface of the hydrophilic and membrane domains. In particular, the process strongly affects the TM5-6^H^ loop, as revealed by our cryo-EM structure of the Y156F^H^/G301L^H^ variant. These findings are consistent with previous studies, biochemically supporting that the loop dynamics is essential for the proton pumping in Complex I^21^. In this regard, Cabrera-Orefice *et al*.^44^ showed that conformationally restraining the nearby TM1-2 loop of NuoA (TM1-2^A^) by introducing a disulphide bridge, resulted in a reduced proton pumping activity, while molecular simulations^21^ and structural^11,12,21,22,45^ studies further identified a conformational coupling network, extending from the β1-β2^NuoCD^ *via* TM1-2^A^ and TM5-6^H^ loops to the TM3^J^ helix. Coupled changes along these key loops could control the conformation of Glu157^H^, and favour the proton transfer along the distal E-channel towards the interface of NuoN, effectively connecting the redox site with the proton pumping modules^12,20,21^. In this regard, we speculate that the rather low activity of the H208A^H^ variant could arise both from the removal of a local proton mediator (HisH^+^) within the E-channel, and from the perturbed hydrogen-bonding network stabilising the *inward* conformation of Tyr156^H^, and thus the coupled movement of Glu157^H^. We note that subtle conformational changes in sidechain position of residues within the E-channel, as observed in the structures of our variants, further support allosteric coupling effects within the pathway.

Based on our combined findings, we suggest that quinone reduction results in proton transfer from Tyr277^CD^ and His228^CD^, which triggers conformational changes in the β1-β2^CD^ loop, the TM1-2^A^ loop, and TM5-6^H^ loop (Fig. 5)^21^. The shifted loop conformations could mediate the conformational flip of Tyr156^H^ from its *outward* conformation (in the *resting* state) to its *inward* conformation (in the *active* state), and mechanically facilitate the π→α transition of TM3^J^. The conformational changes within NuoCD, triggered by the redox reaction, could lead to diffusion of the quinol to site 2, where it forms interactions with Arg87^B^ and the carboxylate cluster of TM5-6^H^, as observed in our MD simulations (Fig. 2a, Supplementary Fig. 3). The *outward* → *inward* flip of Tyr157^H^ positions Glu157^H^ closer to Asp79^A^, thereby favouring proton transfer along the distal part of the E-channel to Glu36^K^/Glu72^K^ pair (*cf*. also Ref^21,33^). Charge accumulation at the interface of NuoN could then induce a conformational change in the conserved Lys217^N^/Glu133^N^ ion-pair of NuoN, favouring lateral proton transfer towards the interface of NuoM, which subsequently results in the propagation of the “charge wave” across NuoM and NuoL. Proton ejection from the open P-side channel of NuoL could result in re-protonation of the hydrophilic axis of NuoL, leading to closing of the ion-pair at the NuoL/M interface, and proton ejection from NuoM. By analogy, the proton uptake and ion-pair closure in NuoM induce proton release from NuoN, while re-protonation and ion-pair closure in NuoN would further eject the proton stored at the end of the E-channel across the membrane. Finally, as a response of the charge redistribution in the E-channel, TM3^J^ returns the π-conformation and the TM5-6^H^ loop relaxes to its resting form, mediated by the Tyr156^H^ backflip to the *outward* conformation. The relaxed loop conformations could, shift the p*K*_a_ of the carboxylate cluster at the Q site 2, favouring re-protonation of QH^-^ and release of a neutral quinol species into the membrane.

**Fig. 5.**
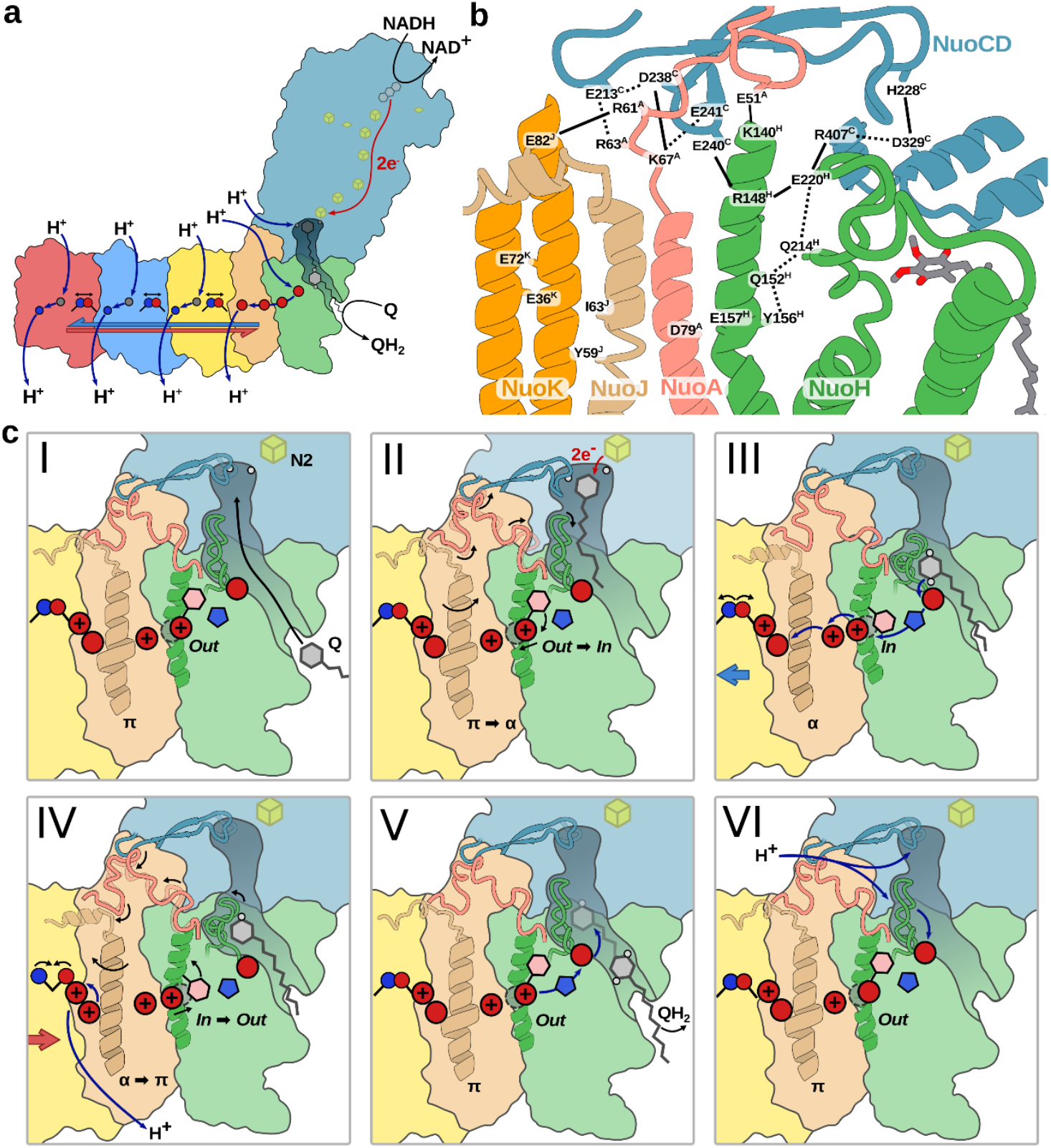
Mechanistic model of redox-driven proton pumping and the involvement of the E-channel. **a**, Schematic overview of the proposed pumping mechanism. Reduction of quinone and binding in Q site 2 initiates an electrostatic wave that propagates along the membrane domain. The protonation wave sequentially induces conformational changes in conserved ion pairs within each antiporter-like subunit, resulting in proton translocation and a single proton being pumped by each subunit. **b**, Overview of the proposed conformational switching network, coupling the Q reduction and movement with the rotation of TM3^J^. Conserved charged and polar residues form a relay, linking the quinol formation and diffusion along the tunnel with structural rearrangements in key functional loops, ultimately activating the proton pumping. **c**, Mechanistic proposal of the conformational switching. Titratable residues are shown as circles, with a “+” denoting the location of protons in the E-channel. Tyr156^H^ and His208^H^ are shown as a hexagon and pentagon, respectively, while the alternative conformation of Glu157^H^ is shown as a grey dashed circle. Black and blue arrows indicate conformational changes and proton transfer processes, respectively. **I:** Q diffuses into the active site through the Q cavity. **II:** reduction of Q to QH_2_ and the coupled QH_2_ diffusion induces conformational changes in the switching network, resulting in conformational changes in the TM1-2^A^, TM5-6^H^, PSST^B^, and β1-β2^CD^ loops. This causes the flip of Tyr156^H^ (*outward* → *inward*) and the rotation of TM3^J^ (π→α). The *inward* Tyr156^H^ shifts the position of Glu157^H^ towards Asp79^A^. **III:** A carboxylate residue on TM5-6^H^ deprotonates QH_2_ at Q site 2, initiating a cascade of proton transfer reactions in the E-channel towards the proton pumping modules. The α-state of TM3^J^ and the coupled conformational changes within the E-channel favour proton transfer from Glu157^H^ and Asp79^A^ towards Glu36^K^, triggering the opening of the ion-pair in NuoN, and initiating the proton pumping process. **IV**: Back-propagation of the protonation wave results in the ion-pair closing, while TM3^J^ rotates to the π state and Tyr156^H^ flips to the *outward* state. The final proton at Glu36^K^/Glu72^K^ is released across the membrane towards the P-side. **V:** The QH^-^ species is re-protonated, triggering the release of QH_2_ into the membrane. **VI:** The residues in the active site and E-channel carboxylate residues are re-protonated from the N-side bulk.

In contrast to our model, in which Tyr156^H^ mechanically mediates conformational coupling between the Q site and the E-channel, Kravchuk *et al*.^12^ suggested that the *inward* flip of Tyr156^H^ enables proton transfer from the E-channel to the quinone substrate bound in the active site in NuoCD upon its 2e^-^-reduction. In their “NuoL-only” pumping model, where only the terminal subunit is suggested to pump all protons, the resting state of Complex I comprises an *outward* Tyr156^H^ and was suggested to result in a dry E-channel blocking the proton transfer reaction from the E-channel to the Q site. Recently a computational study^46^ performed on the mouse Complex I without significant structural relaxation (but *cf*. also Ref^20,21^), supported the proposal of Kravchuk *et al*. that the Tyr156^H^ flip is critical for the proton transfer reaction itself, thus contradicting our current results. Moreover, the authors suggested that the phenolic group cannot participate in the proton transfer reaction, in contrast to our large-scale QM/MM free energy explorations, where the phenol mediates proton exchange with low free energy barriers. Although the origin of these discrepancies is unclear, these observations could arise from the highly strained geometric reaction coordinate applied on specific bonds in their calculations resulting in a possible bias on the explored reactions, together with large drifts and statical error in the resulting free energy profiles. In contrast, we have adopted here the modified centre of excess charge (mCEC), which allows a broad exploration of the free energy landscape on long QM/MM-MD timescales together with (classical) dynamic relaxation of key intermediates to enhance the convergence.

The proton transfer reactions along the E-channel was recently also addressed by multi-state empirical valence bond (MS-EVB) simulations^47^, suggesting that the reaction is mediated only by water molecules and carboxylates, with the proton transfer resulting in a drastic hydration shift, leading to an influx of over 40 water molecules between Glu157^H^, Asp79^A^, and Glu36^K^/Glu72^K^ alone. However, due to limitations in the employed MS-EVB model, the authors did not model proton transfer reactions between any other protein residues besides carboxylates, which could explain, in part, the different molecular principles from those observed here and in Ref^33^. Importantly, our DFT-based QM/MM simulations do not rely on a pre-parametrized force field, but treat the protonation reactions of *all* amino acids on an equal footing, based on first-principles calculations. Moreover, even if longer simulation timescales of the EVB simulations can enhance the free energy convergence, we speculate that the relaxation of a hydronium species at transient sites may cause artificial flooding effects similar to those observed when all carboxylates of the E-channel are modelled in their fully charged (anionic) state. Nevertheless, while this study generally supports our findings on the central role of water molecules in enabling the proton transfer reactions in Complex I (cf. also Ref. ^18,21,45,46,48^) and other bioenergetics protein complexes^38,43^, we note that the reported drastic water influx and resulting structural changes in Ref.^47^ is not supported by current cryo-EM data or our multiscale simulations, and does not account for the Tyr156^H^ or His208^H^-mediated proton transfer reactions described here.

Although our calculations do not directly assess the directionality of the proton transfer along the E-channel, our MD simulation suggest that a neutral Glu157^H^ and a deprotonated Glu216^H^, which is a prerequisite configuration for proton transfer towards the quinone cavity (Glu157^H^ → Glu216^H^) in the Kravchuk *et al*. model, lacks a water-mediated connection between the Glu157^H^ and Tyr156^H^ or His208^H^ (Supplementary Fig. 4j), suggesting that proton transfer in this direction and configuration is unlikely.

Importantly, although we find that Tyr156^H^ *can* mediate proton exchange, our data show that the Tyr156^H^ is not essential for the proton transfer reaction within the E-channel, as the exchange of the phenolic group with a phenyl moiety does not block the process. Taken together, we suggest that Tyr156^H^ functions as a mechanical coupling site, mediating long-range conformational crosstalk across an intricate structural network.

## Conclusions

By combining multiscale simulations with site-directed mutagenesis, proteoliposome experiments, and structural analysis, we have addressed here the functional role of Tyr156^H^ in mediating the redox-driven proton pumping process in complex I. We showed that the Tyr156^H^ flip, which is conserved across Complex I from different species, is mechanically coupled to the rotation of TM3^J^ with conformational changes in surrounding conserved loops (TM5-6^H^ and TM1-2^A^). These coupled rearrangements are likely to trigger the redox-driven proton pumping process in the membrane domain of the complex. Our *in silico* and *in vitro* mutagenesis experiments show that Tyr156^H^ is not essential for the proton transfer itself, but could instead function as a molecular latch that regulates the directionality of the proton transfer process, during forward and reverse conditions. Together, our findings reveal that the strong electrodynamic coupling principles driving the long-range proton pumping process in Complex I, also rely on protein dynamics mediated by conserved helices and loops. On a general level, our findings illustrate the intricate interplay between protein dynamics and catalysis in mediating long-range *action-at-a-distance* effects.

## Methods

### Molecular dynamics simulations

Classical MD simulations were constructed based on the cryo-EM structure of *E. coli* Complex I (PDB ID: 7Z7S^12^), and previous work^49^. The protein was embedded in a lipid bilayer comprising 70% POPE, 20% POPG, and 10% cardiolipin, solvated in TIP3P water box, and neutralized with 150 mM NaCl (Supplementary Fig. 1). The FeS clusters and FMN were modelled in their oxidized state, while quinol or a QH^-^ species was modelled in the substrate tunnel at the Q site 2 based on previous MD simulations of *Y. lipolytica*^21^ and *M. musulus*^20^ Complex I. Protonation states were assigned based on prior p*K*_a_ calculations^49^, with Glu218^H^ and Glu241^H^ modelled in their protonated forms, His208^H^ was modelled in its ε-protonated form, while other acidic residues along the E-channel were deprotonated, unless otherwise stated (Supplementary Table 3). The simulation system comprised ∼853,000 atoms. MD simulations were run in an *NPT* ensemble at 310 K and 1 atm using a 2 fs timestep. Long-range electrostatics were treated using the particle mesh Ewald (PME) method, with a grid size of 1 Å. The system was treated using the CHARMM36 force field^50^, whereas force field parameters for quinol and FeS clusters based on in-house DFT parametrization. *In silico* mutations were introduced *via* psfgen module in VMD, followed by 2 ps equilibration at 310 K. Each system was energy minimized, followed by heating from 0 K to 310 K in the *NVT* ensemble, and subsequent equilibration under *NPT* conditions at 1 atm using a Langevin thermostat and barostat. All MD simulations were performed using NAMD (versions 2.14 and 3.0)^51^, and analysed using VMD^52^ and MDAnalysis^53,54^. The proton pathways were characterised using the water bridge analysis module in MDAnalysis^53,54^. Electric field strength along the proton pathway was computed by projecting local electric field vectors calculated with TUPA^55^ onto the local tunnel axis of the CAVER^56^-identified pathway. See Supplementary Fig. 1 for system setup, and Supplementary Tables 1 and 3 for details of the MD simulations.

### Classical free energy simulations

Umbrella sampling (US) simulations were performed to explore the *inward* → *outward* flip of Tyr156^H^ (Phe156^H^). Initial windows for the US simulations were generated based on steered MD (SMD) simulations, performed on the equilibrated, unbiased MD trajectories. The SMD simulations were carried out by biasing the dihedral angle between backbone atoms (C, N) and sidechain atoms (Cγ, C?) with a harmonic restraint of 0.2 kcal mol^-1^ deg^-2^ from -80° to 72° over 20 ns (Supplementary Fig. 8), as implemented in the Colvars module of NAMD 3.0.^51^ US simulations were performed based on relaxed structures extracted from the SMD trajectory. US simulations were carried out in 39 windows harmonic restraint of 0.2 kcal mol^-1^ deg^-2^, resulting in a total sampling of 390-585 ns for each free energy profile, derived by WHAM/MBAR, while statistical errors were computed based on bootstrap analysis. See Supplementary Fig. 7 and Supplementary Table 3 for further details.

### QM/MM free energy simulations

Hybrid QM/MM simulations were performed for exploration of the proton transfer reactions in the E-channel, based on representative structural snapshots extracted from classical MD simulations. The QM/MM system was truncated to ∼12,000-16,400 atoms, by retaining all atoms within 23 Å of the QM region. Residues within 10 Å of the QM region were allowed to move during the QM/MM simulations, while atom beyond this range were fixed during the simulations (Supplementary Fig. 1). The QM region (see Supplementary Table 2), with *N*=121-156 atoms, was treated at the B3LYP-D3/def2-SVP level^57–60^, and coupled to MM system using an additive electrostatic embedding scheme with the link atom approach, introduced between Cα-Cβ atoms. The classical region was modelled using the CHARMM36 force field^50^ in combination with in-house parameters (see above). All systems were gradually heated from 0.1 K to 310 K prior to sampling for equilibration. The QM/MM simulations were carried out using FermiONs++^61^ and OpenMM^62^. See Supplementary Fig. 5 and Supplementary Table 2 for details of the reaction pathways and QM region.

The QM/MM free energy profiles for all proton transfer reactions were modelled using the modified centre of excess charge (mCEC)^63,64^ as a reaction coordinate using umbrella sampling (US). Each US window was initiated sequentially from the final coordinates of the preceding window to ensure continuity of sampling along the reaction pathway, followed by 4-6 ps sampling of each window along the mCEC coordinate, with harmonic restraints, using a force constant of 100 kcal mol^-1^ Å^-2^ and a timestep of 0.5 fs, and with a separation between consecutive windows of 0.25 Å on the reaction coordinate. The total QM/MM free energy sampling timescale was 1012 ps (see Supplementary Table 2), but performed on classically relaxed structures, which account for dynamic effects on the μs-timescales.^65^

### Protein production and purification

#### Cell growth

*E. coli* strain BW25113 containing the pBAD_nuo_ plasmid, was cultivated in either M9 minimal medium supplemented with additives (1:100, *v/v*) or in LB medium, each containing the relevant antibiotics when needed (see Supplementary Table 6)^49^. Pre-cultures were inoculated from a liquid culture, glycerol stock, or single colony, and grown at 37°C with shaking at 180 rpm. Day cultures were inoculated 1:100 (v/v), while overnight cultures were inoculated 1:1000 (v/v). For protein production, 400–800 mL of autoinduction medium was inoculated 1:13.33 (v/v) with pre-culture and incubated at 37°C with shaking at 180 rpm. Cell growth was monitored by measuring optical density at 600 nm (OD_600_). Cells were harvested at OD_600_ ≈ 4.0 by centrifugation at 5749 g for 20 min at 4°C (Rotor JLA-81000, Avanti J-26 XP, Beckman Coulter), flash-frozen in liquid nitrogen, and stored at -80°C until further use.

#### Isolation of cytoplasmic membranes

All steps were performed at 4°C or on ice unless otherwise stated. *E. coli* cells (6-50 g) were suspended in five volumes ice-cold A buffer (Supplementary Table 6) supplemented with 1 mM PMSF and DNase I. Cells were homogenized using a Teflon–glass homogenizer and broken *via* high-pressure homogenization (HPL6, Maximator; 3 cycles, 1000–1500 bar). The suspension was clarified by centrifugation at 12,074 *g* for 20 min at 4°C to remove unbroken cells and debris. The supernatant was ultracentrifuged at 178,000 *g* for 70 min (Rotor 60Ti, Beckman) to sediment the cytoplasmic membranes. The sediment was resuspended in A* buffer (pH 6.8, 1:2 w/v) containing PMSF (1:1000, v/v) using a Teflon–glass homogenizer, flash-frozen in liquid nitrogen, and stored at −80°C until further use.

#### Purification of Complex I and variants

All following steps were carried out at 4°C or on ice. Buffers contained 30 μM PMSF unless otherwise noted. Cytoplasmic membranes (50 g) were thawed and diluted 1:3 (w/v) in A* buffer (pH 6.8, Supplementary Table 5). Membrane proteins were extracted by adding LMNG (5%, w/v stock) to a final concentration of 2%, followed by gentle stirring for 1 h at room temperature. The extract was diluted with 100 mL A* buffer, clarified by ultracentrifugation (178,000 g, 15 min, Rotor 60Ti, Beckman Coulter), and filtered through a 0.45 μm membrane (Filtropur S 0.45, Sarstedt).

For affinity chromatography, the clarified extract was adjusted to 20 mM imidazole and applied to a 35 mL ProBond Ni^2+^-IDA column (BioRad) equilibrated with binding buffer. The column was washed with buffer containing 116 mM imidazole (20% elution buffer) and eluted with 308 mM imidazole (60% elution buffer) at 2 mL min^-1^. The NADH/ferricyanide oxidoreductase activity of the 4 mL fractions was assayed, active fractions were pooled and concentrated to 2 mL using a 100 kDa MWCO spin concentrator (Amicon Ultra-15, Millipore), followed by clarification at 17,500 *g* for 10 min (Eppendorf 5417 R).

The concentrate was loaded at 0.3 mL min^-1^ onto a 297 mL Superose 6 size-exclusion column equilibrated in A* buffer containing LMNG. 2 mL fractions were collected between 80 mL and 250 mL elution volume. The most active fractions were combined and concentrated to 20 mg mL^-1^. Purified Complex I was snap-frozen in liquid nitrogen and stored at -80°C. See Supplementary Fig. 10 for chromatograms of protein purification and SDS-PAGE analysis.

#### Site-directed mutagenesis

DNA fragments were amplified using a LifeTouch thermocycler (Bioer) in 10 μL or 50 μL reaction volumes (0.2 mL tubes, Biozym). PCR mixtures were prepared on ice using double-distilled water (ddH_2_O) and screened with and without 5% (v/v) DMSO for optimal annealing temperatures. Products from 10 μL reactions were analysed by agarose gel electrophoresis and the optimal conditions were used for 50 μL scale-up reactions.

PCR products were treated with DpnI to digest methylated parent DNA. Site-directed mutagenesis was performed following the QuikChange protocol (Agilent Technologies) using 50 μL reactions containing 70– 100 ng template DNA, 0.2 μM oligonucleotides, 20 μM dNTPs, 1× KOD Hot Start buffer, 2.25 mM MgSO_4_, and 0.2 U μL^-1^ KOD Hot Start DNA polymerase. DMSO (5%, v/v) was added when necessary to facilitate amplification of GC-rich or structurally complex templates. Thermal cycling conditions followed the manufacturer’s instructions. Successful amplification and correct insert size were confirmed by agarose gel electrophoresis. Designed primers and used plasmids are listed in Supplementary Table 7.

#### Preparation of proteoliposomes

Proteoliposomes were prepared by reconstituting purified *E. coli* Complex I into preformed liposomes composed of *E. coli* polar lipids (Avanti). Lipid stocks (1 mL in chloroform) were dried under a nitrogen stream and resuspended in five volumes of liposome buffer. The lipid suspension was frozen in liquid nitrogen and thawed at 29°C in a thermomixer (Thermomixer Compact, Eppendorf) seven times to promote unilamellar vesicle formation. Vesicle size was homogenized by extruding 250 μL liposome suspension ≥21 times through a 0.1 μm polycarbonate membrane (Mini Extruder, Avanti). Extruded vesicles were supplemented with 8 μL 20% (w/v) sodium cholate to facilitate membrane protein incorporation.

17 μL reconstitution buffer (Supplementary Table 6) were added to 50 μL Complex I (20 mg mL^-1^), incubated on ice for 5 min and combined with 250 μL liposome suspension. After 20 min incubation at room temperature, detergent was removed by applying the mixture to a PD-10 desalting column (8.3 mL; Sephadex G-25, GE Healthcare) equilibrated in liposome buffer. The flow-through was collected and centrifuged at 150,000 *g* for 30 min at 4°C (Rotor A-100, Airfuge, Beckman) under 2 bar air pressure. Sedimented proteoliposomes were resuspended in 500 μL liposome buffer and used immediately to determine enzymatic activities (see below).

The orientation and reconstitution efficiency of Complex I were assessed by measuring NADH/ferricyanide oxidoreductase activity. Since NADH cannot cross intact lipid bilayers only *outward*-facing Complex I contributes to this activity. To determine total Complex I content, proteoliposomes were dissolved with 0.04% (w/v) DDM, making all Complex I accessible to NADH. The ratio of activity before and after detergent treatment was used to calculate the proportion of the two possible enzyme orientations.

#### NADH/ferricyanide oxidoreductase activity

The NADH:ferricyanide oxidoreductase activity of Complex I and variants was quantified spectrophotometrically by monitoring the decrease in absorbance of potassium ferricyanide (K_3_[Fe(CN)_6_]) at 410 nm (ε_410_ = 1 mM^−1^ cm^−1^). Reactions were carried out in 1 mL A* buffer (Supplementary Table 5) containing 1 mM K_3_[Fe(CN)_6_] and 0.2 mM NADH. The reaction was initiated by the addition of 0.1–50 μL protein solution. Measurements were performed using either an Ultrospec 1000 Pro spectrophotometer (Amersham Bioscience) or a diode-array spectrophotometer (TIDAS S, J&M Analytik AG) equipped with 1 cm pathlength UV cuvettes (Macro QS, Hellma). To distinguish enzymatic from non-enzymatic electron transfer activity, absorbance changes were recorded for 30 s before and after the addition of the sample. The rate of Complex I activity was determined by subtracting the background signal from the total absorbance change.

#### NADH:decylubiquinone oxidoreductase activity assay

The physiological NADH:decylubiquinone oxidoreductase activity of Complex I and variants were assessed by monitoring the decrease in NADH consumption at 340 nm (ε_340_= 6.178 mM^−1^ cm^−1^) using a diode-array spectrophotometer (TIDAS S, J&M Analytik AG; 1 cm pathlength UV cuvettes, Macro QS, Hellma) at 30°C. The assay contained 60 μM decyl-ubiquinone and 20 μL proteoliposomes in 1 mL A* buffer (Supplementary Table 6). After 1 min pre-incubation, the reaction was initiated by adding 150 μM NADH.

#### Measurement of pH-gradient by quenching the ACMA fluorescence

The proton pumping activity of Complex I was assessed in freshly prepared proteoliposomes by monitoring the quenching of the pH-sensitive dye 9-amino-6-chloro-2-methoxyacridine (ACMA). Fluorescence was measured in ACMA buffer (30°C; Supplementary Table 6), containing 100 μM DQ, 2 μM ACMA, and 50 μL proteoliposomes in a final volume of 1 mL. The fluorescence emission at 480 nm (excitation: 430 nm; slit width: 2.5 nm) was recorded using a PerkinElmer LS 55 fluorescence spectrometer (Hellma Macro-Special quartz cuvettes, 1 cm pathlength). After 2 min equilibration, proton translocation was initiated by the addition of 130 μM NADH. Dissipation of the resulting proton gradient (ΔpH) was achieved by the addition of 1 μg mL^-1^ of gramicidin.

#### Measurement of membrane potential by changes of the oxonol IV absorbance

The generation of a membrane potential (ΔΨ) was monitored by changes in oxonol VI absorbance, measured at 588 and 625 nm, respectively. Spectra were recorded using a diode-array spectrophotometer (TIDAS S, J&M Analytik AG) with 1 cm pathlength cuvettes (Macro QS, Hellma) and the kinetic traces at 588 and 625 nm were extracted. The amount of proteoliposomes added to each reaction was normalized based on the proportion of outward-orientated Complex I (see above). Measurements were performed in oxonol buffer (Supplementary Table 6), containing 50 μM DQ 0.5 μM oxonol VI (freshly prepared 50 mM stock in methanol, vortexed thoroughly), and the proteoliposomes. After an initial baseline acquisition of 50 s, the reaction was initiated by the addition of 100 μM NADH. The membrane potential was dissipated after 150 s by addition of gramicidin (1 μg mL^−1^).

#### Cryo-EM grid preparation and data collection

Freshly purified Complex I (∼10 μg μL^-1^, 9.8 μL) was supplemented with 0.2 μL of 5% (w/v) Fos-Choline-10 to improve particle dispersion. The mixture was incubated on ice for 3 min prior to vitrification. Grids (R 2/1 on 300 gold mesh, Quantifoil) were glow-discharged (PELCO easiGlow), and 3 μL of the sample was applied to each grid. Grids containing the variants were vitrified using a Vitrobot Mark IV (Thermo Fisher Scientific) at 100 % humidity, at 4°C and blotted for 6 s with filter paper. The Y156F sample was flash frozen into liquid propane, while the G301L, G301L/Y156F and H208A variants were frozen in liquid ethane. All grids were stored in liquid nitrogen until further use. Data were collected using SerialEM on a 300 kV Titan Krios transmission electron microscope (Thermo Fisher Scientific) equipped with a Gatan K2 summit direct electron detector in counting mode with a pixel size of 0.572 Å px^-1^ or 0.931 Å px^-1^, 40 frames per image and a total dose of 40 e^-^Å^-2^.

#### Data processing, model building and refinement

The micrographs were processed in cryoSPARC v4.7.1. Raw movie stacks were motion corrected and contrast transfer function (CTF) estimated using patch motion correction and patch CTF estimation. Particle picking was done with a template generated from the cryo-EM structure of *E. coli* Complex I (PDB ID: 7Z7S), followed by an extraction from micrographs, exposure group split and two-dimensional (2D) classification to generate 2D classes. After 2D classification of the binned particle set, selected particles were used for generating 3D initial models by *ab initio* reconstruction with multiple classes. The particles from the best class were first heterogeneously and non-uniformly (NU) refined. The datasets were unbinned with a box size of 512 px, 720 px or 800 px.

NU-refinements yielding final refined maps from 99,143 particles at 2.74 Å resolution for the Y156F^H^ variant, from 107,112 particles at 2.57 Å for the Y156F^H^/G301L^H^ variant, from 121,679 particles at 2.99 Å for the G301L variant, from 150,920 particles at 2.31 Å for the H208A^H^ variant. The gold standard Fourier shell correlation (FSC) 0.143 criteria were used to provide the map resolution estimate. Composite maps were created from each of the last NU-refinements and focused maps (peripheral arm, membrane arm, junction) of the variants. These were created by particle subtraction, followed by local refinement using the corresponding mask. These procedures yielded higher resolutions of 2.37–2.54 Å for the Y156F variant, 2.28–2.40 Å for the Y156F^H^/G301L^H^ variant, 2.68–2.83 Å for the G301L^H^, 2.11–2.16 Å for the H208A^H^ variant. Details of the workflow of the structure solution are provided in Supplementary Figs. 12-15. The resulting volumes were merged in ChimeraX. The cryo-EM structures of *E. coli* Complex I (PDB IDs: 7Z7S and 9Q8I) were used as initial models for the variants, fitted into the merged maps using ChimeraX and manually corrected and built in COOT. Final refinement and the extraction of data and model statistics were done in PHENIX. The cryo-EM data analysis and validation are shown in Supplementary Table 5 and Supplementary Figs. 12-16. Example densities are shown in Supplementary Fig. 16.

Atomic model building was performed in Coot (v0.9.7)^66^ using the cryo-EM density maps as a guide. Iterative real-space refinement and validation were carried out in Phenix^67,68^, followed by manual inspection and adjustment to ensure consistency with the experimental density and stereochemical restraints.

## Acknowledgments

We thank Dr. Terezia Kovalova and Dr. Daniel Wohlwend for technical help with cryo-EM models and Zoé Kupek for help with the protein characterisation. This work was supported by the Knut and Alice Wallenberg Foundation (2019.0251, 2024.0220, and WASPDDLS22:025 to V.R.I.K), the Göran Gustafsson Foundation for Research in Natural Sciences and Medicine (to V.R.I.K.), by the Swedish Research Council (2020-04081 and 2025-04607 to V.R.I.K.), and by the Deutsche Forschungsgemeinschaft (DFG GRK2202; 235777276/RTG & SPP1927; FR 1140/11-2 to T.F.). Computing resources were provided by the National Academic Infrastructure for Supercomputing in Sweden (NAISS 2025/1-33, 2024/1-28). We thank Dr. Stefan Steimle for technical help in collecting the cryo-EM data at the cryo-EM Facility of the University of Freiburg, with the Titan Krios G4, operated within the Microscopy and Image Analysis Platform (MIAP), University Freiburg, funded by the Deutsche Forschungsgemeinschaft (project nr. 506518771). We also thank cryo-EM Swedish National Facility funded by the Knut and Alice Wallenberg, Family Erling Persson and Kempe Foundations, SciLifeLab, Stockholm University and Umeå University for supporting our work.

## Data availability

Data supporting the findings of this manuscript are available from the corresponding authors upon reasonable request. A reporting summary for this Article is available as a Supplementary Information file. The source data underlying Figs. 1-5 and Supplementary Figs. 1-18 are provided as a Source Data file. Model and cryo-EM density maps of the Y156F^H^, G301L^H^, Y156F^H^/G301L^H^ and H208A^H^ variant Complex I were deposited to the PDB under accession codes 28WM, 28VI, 28TL, 28NI and the EM Data Bank under accession codes EMD-56915, EMD-56916, EMD-56917, EMD-56918, EMD-56913 for Y156F^H^, EMD-56880, EMD-56881, EMD-56882, EMD-56883, EMD-56884 for G301L^H^ and EMD-56649 EMD-56653, EMD-56654, EMD-56655, EMD-56656 for H208A^H^. MD simulation setups and structural snapshots are available in the following Zenodo repository: 10.5281/zenodo.18847236.

## Author contributions

V.R.I.K. and T.F. designed the study and directed the project;

N.L.H., J.H, and Th.S. constructed, isolated, and experimentally characterized the protein;

A.B. and Ti.S. assisted in the experiments;

N.L.H., performed multiscale simulations

N.L.H., J.H., and Th.S. collected cryo-EM data;

N.L.H., J.H., and Th.S. refined cryo-EM models;

N.L.H., J.H., H.K., Th.S., T.F., and V.R.I.K. analysed cryo-EM data

N.L.H., J.H., H.K., Th.S., A.B., Ti.S., P.S., T.F., V.R.I.K. analysed data

N.L.H., H.K., T.F., V.R.I.K. wrote the manuscript with input from all authors.

## Competing interests

The authors declare no competing interests.

## Additional information

Supplementary information is available for this paper at XXX.

Correspondence and requests for materials should be addressed to T.F. and V.R.I.K.

## Notes

### Competing Interest Statement

The authors have declared no competing interest.

