## Supplementary Information for "Redox-Triggered Coupling Network Mediates Long-Range Energy Transduction in Respiratory Complex I"

### Content

**Supplementary Fig. 1 |** MD and QM/MM setups.
**Supplementary Fig. 2 |** Central distances in the E-channel from MD simulations.
**Supplementary Fig. 3 |** Central distances in Q site 2 from MD simulations.
**Supplementary Fig. 4 |** Hydrogen-bonded connectivity along the E-channel.
**Supplementary Fig. 5 |** Convergence of QM/MM free energy simulations, and structure of intermediates along the proton transfer process.
**Supplementary Fig. 6** | Electric field effects along the proton pathway in the WT and Complex I variants.
**Supplementary Fig. 7** | Free energy profile of the *inward* → *outward* flip of Tyr156$^{H}$.
**Supplementary Fig. 8 |** Comparison of the *inward*/*outward* conformation of Tyr156$^{H}$ in cryo-EM structures of Complex I.
**Supplementary Fig. 9** | Structural re-arrangements from TMD simulation.
**Supplementary Fig. 10** | Chromatographic profiles for protein purification and SDS-PAGE of all preparations.
**Supplementary Fig. 11** | Proton translocation and membrane potential formation in the NuoH variants.
**Supplementary Fig. 12** | Cryo-EM data analysis and validation of the 2.7 Å map of the Y157F$^{H}$ variant.
**Supplementary Fig. 13** | Cryo-EM data analysis and validation of the 2.9 Å map of the Y157F$^{H}$/G301K$^{H}$ variant.
**Supplementary Fig. 14** | Cryo-EM data analysis and validation of the 2.9 Å map of the G301K$^{H}$ variant.
**Supplementary Fig. 15** | Cryo-EM data analysis and validation of the 2.2 Å map of the H208A$^{H}$ variant.
**Supplementary Fig. 16** | Example densities based on the 2.2-2.9 Å maps of the variants.
**Supplementary Fig. 17** | Structure of different conformational states of Tyr156$^{H}$ and Glu157$^{H}$ in various NuoH variants.
**Supplementary Fig. 18** | Comparison of RMSF form MD simulations with beta factors form cryo-EM data.

**Supplementary Table 1 |** List of performed MD simulations and modelled states.
**Supplementary Table 2 |** List of performed QM/MM free energy simulations.
**Supplementary Table 3 |** List of performed classical free energy simulations.
**Supplementary Table 4 |** List of base protonation states used in the MD simulations.
**Supplementary Table 5 |** Cryo-EM data collection, refinement, and validation statistics.
**Supplementary Table 6 |** List of buffer compositions.
**Supplementary Table 7 |** Designed primers and plasmids.
**Supplementary Table 8** | NADH:DQ oxidase activity of *E. coli* Complex I (WT) and variants.

**Supplementary Movie 1 |** Proton transfer along the E-channel and the Tyr156$^{H}$ flip.
**Supplementary Movie 2 |** MD simulations of the conformational flip of Tyr156$^{H}$.

**SI References**

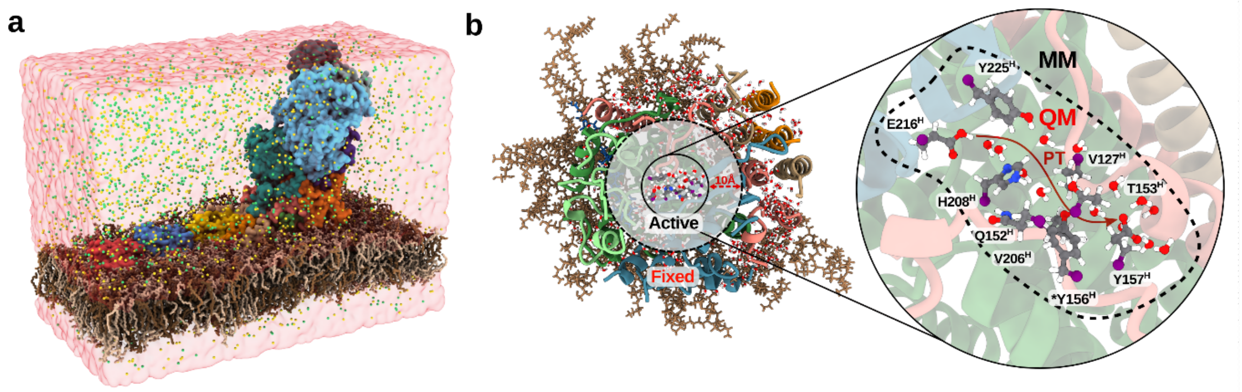


**Supplementary Fig. 1 | MD and QM/MM setups. a,** Overview of the MD simulation system of *E. coli* Complex I, embedded in a lipid bilayer and solvated in an explicit water box with sodium chloride (yellow, green) ions. **b,** QM/MM model used for free energy simulations of the proton transfer along the E-channel.

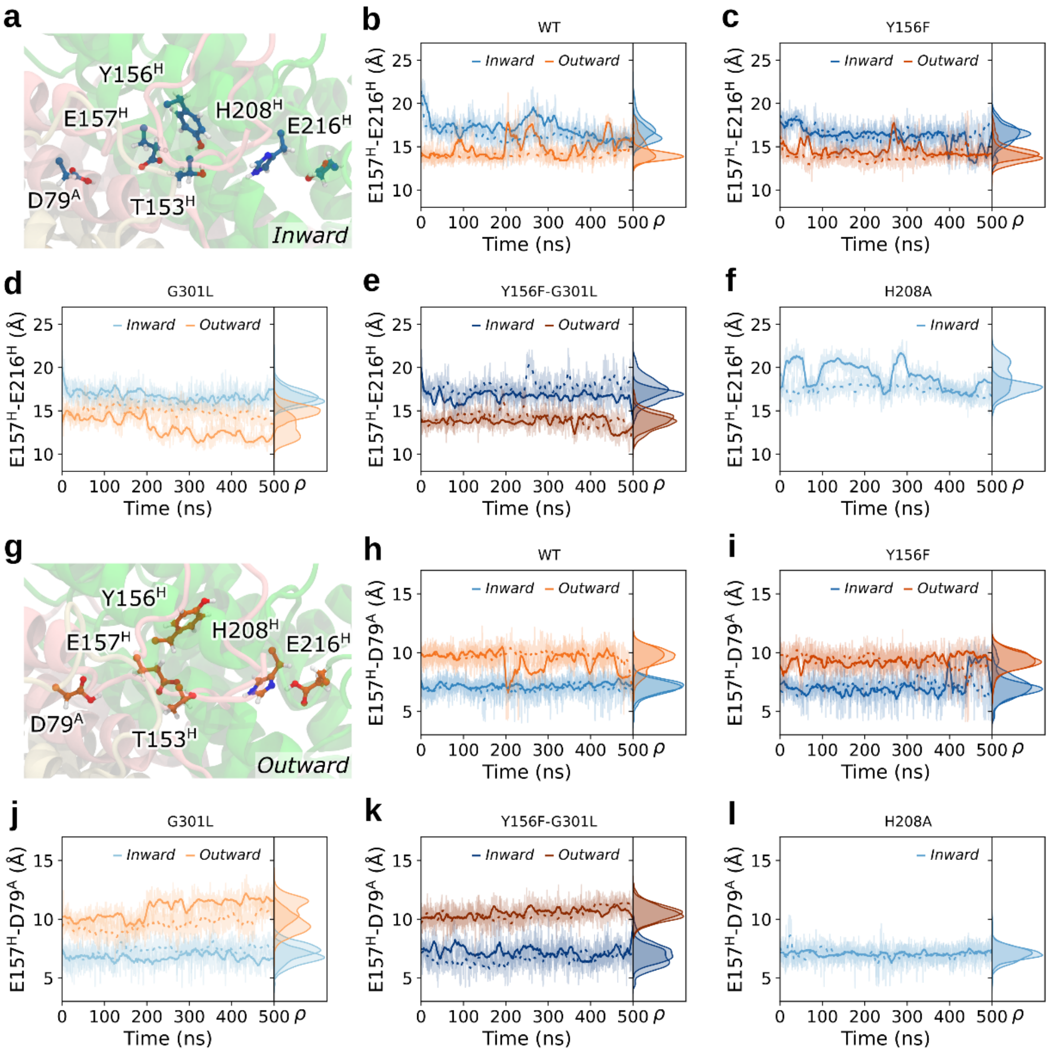


**Supplementary Fig. 2 | Central distances in the E-channel from MD simulations. a** and **g,** Snapshots from MD simulations with Tyr156$^{\mathrm{H}}$ in the **a,** *inward* and **g,** *outward* conformations. **b-f,** The distance between Glu216$^{\mathrm{H}}$ and Glu157$^{\mathrm{H}}$ from **b,** WT; **c,** Y156F$^{\mathrm{H}}$; **d,** G301L$^{\mathrm{H}}$; **e,** Y156F$^{\mathrm{H}}$/G301L$^{\mathrm{H}}$; and **f,** H208A$^{\mathrm{H}}$ simulations. The simulation replicas are shown as dotted lines. **h-l,** The distance between Glu157$^{\mathrm{H}}$ and Asp79$^{\mathrm{A}}$ from **b,** WT; **c,** Y156F$^{\mathrm{H}}$; **d,** G301L$^{\mathrm{H}}$; **e,** Y156F$^{\mathrm{H}}$/G301L$^{\mathrm{H}}$; and **f,** H208A$^{\mathrm{H}}$ simulations.

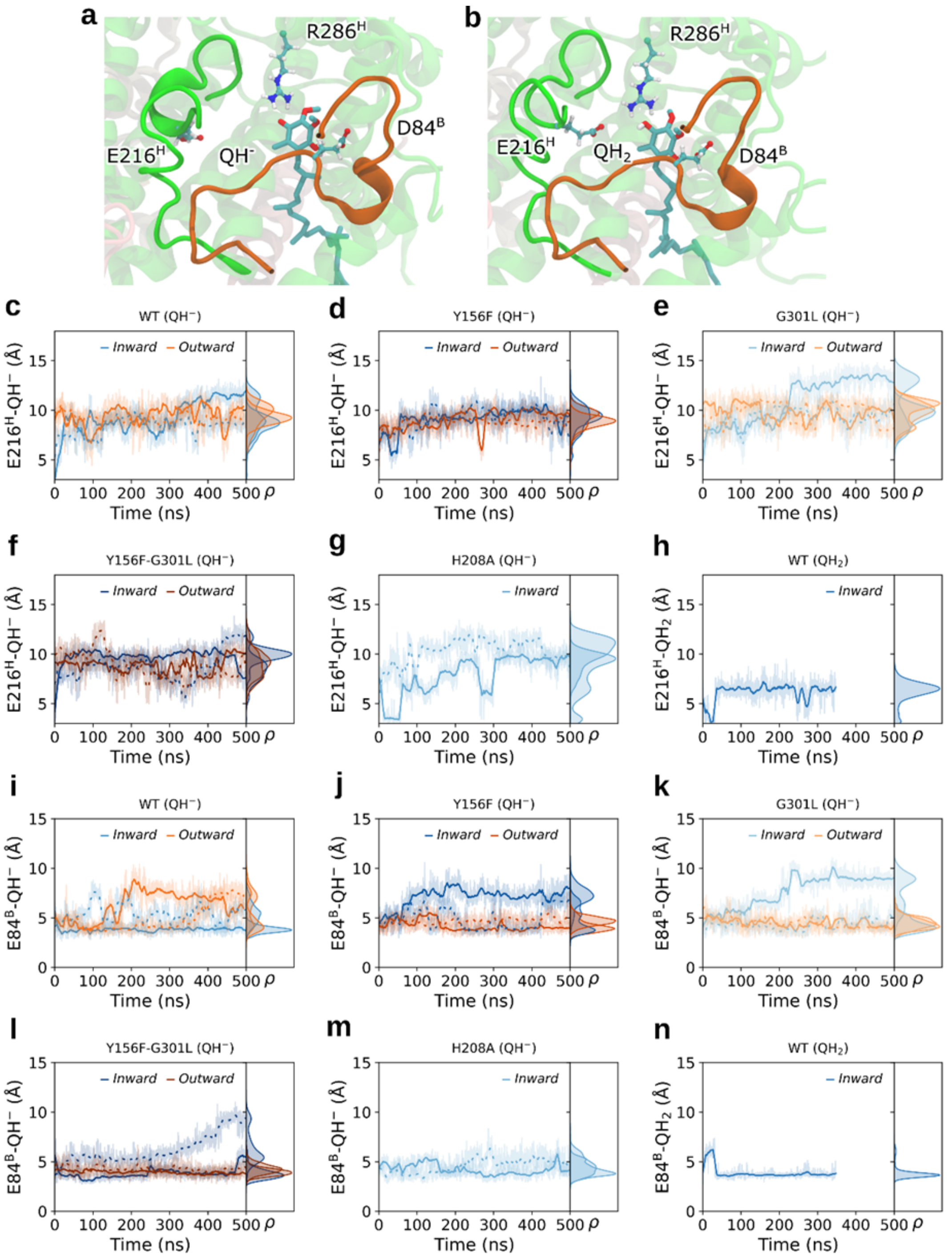


**Supplementary Fig. 3 | Central distances in Q site 2 from MD simulations. a-b,** Representative snapshots from MD simulations showing the **a,** $QH^-$ and **b,** $QH_2$ interactions in the Q site 2. **c-h,** Distance between Glu216$^H$ and $QH^-/QH_2$ from MD simulations with **c,** WT ($QH^-$); **d,** Y156F$^H$; **e,** G301L$^H$; **f,** Y156F$^H$/G301L$^H$; **g,** H208A$^H$; and **h,** WT ($QH_2$). **i-n,** Distance between Glu84$^H$ and $QH^-/QH_2$ from MD simulations with **i,** WT ($QH^-$); **j,** Y156F$^H$; **k,** G301L$^H$; **l,** Y156F/G301L$^H$; **m,** H208A$^H$, and **n,** WT ($QH_2$).

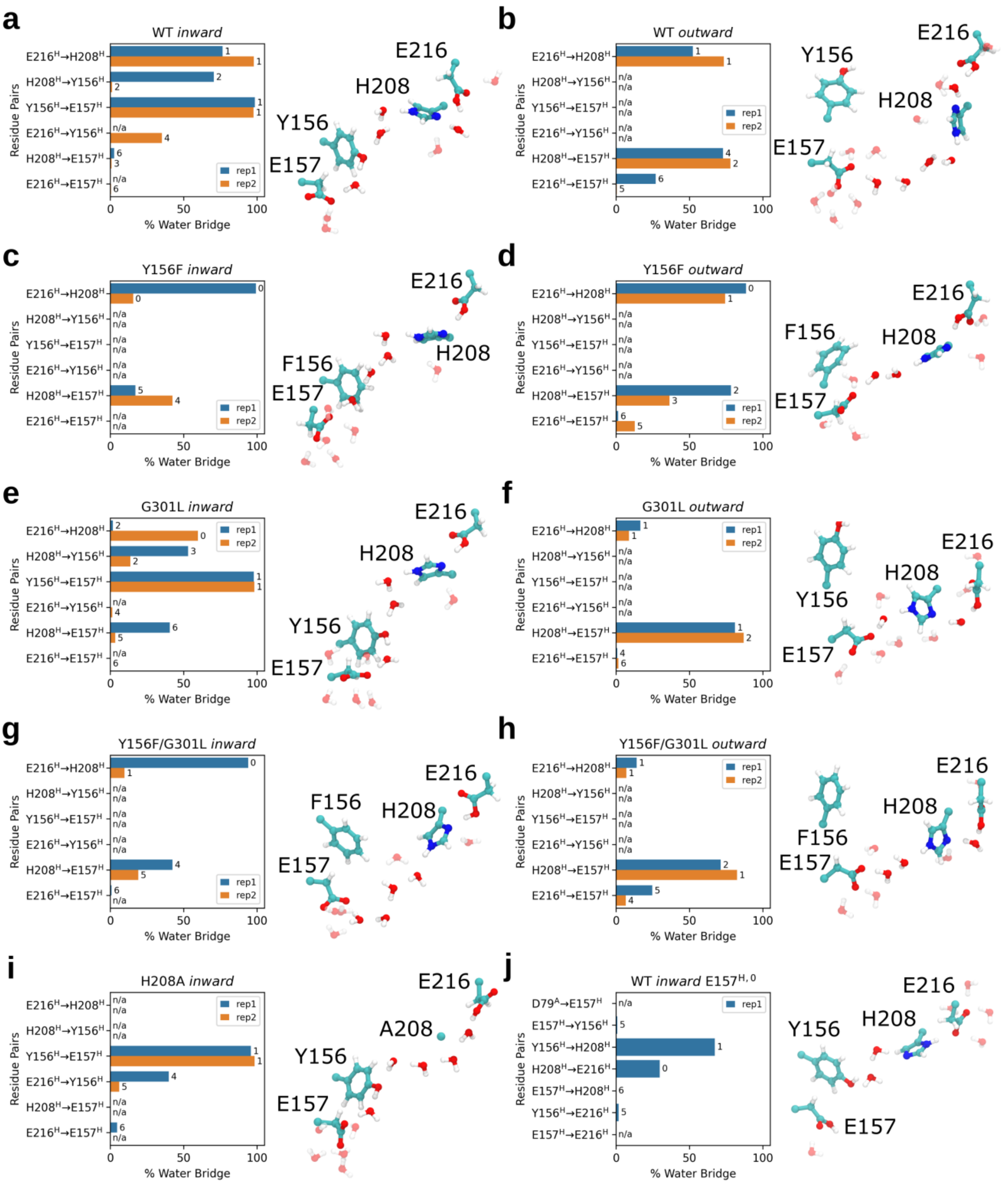


**Supplementary Fig. 4 | Hydrogen-bonded connectivity along the E-channel. a-j,** ***Left***: Percentage of residue pairs in hydrogen-bonded contact during the last 100 ns of the MD simulations of each variant. The directional hydrogen-bonding interactions are mediated by water molecules, with the median water count shown next to each bar. ***Right***: Snapshot from MD simulations showing a representative structure of the hydrogen-bonded network within the cluster. The water molecules directly involved in the wire are shown in opaque.

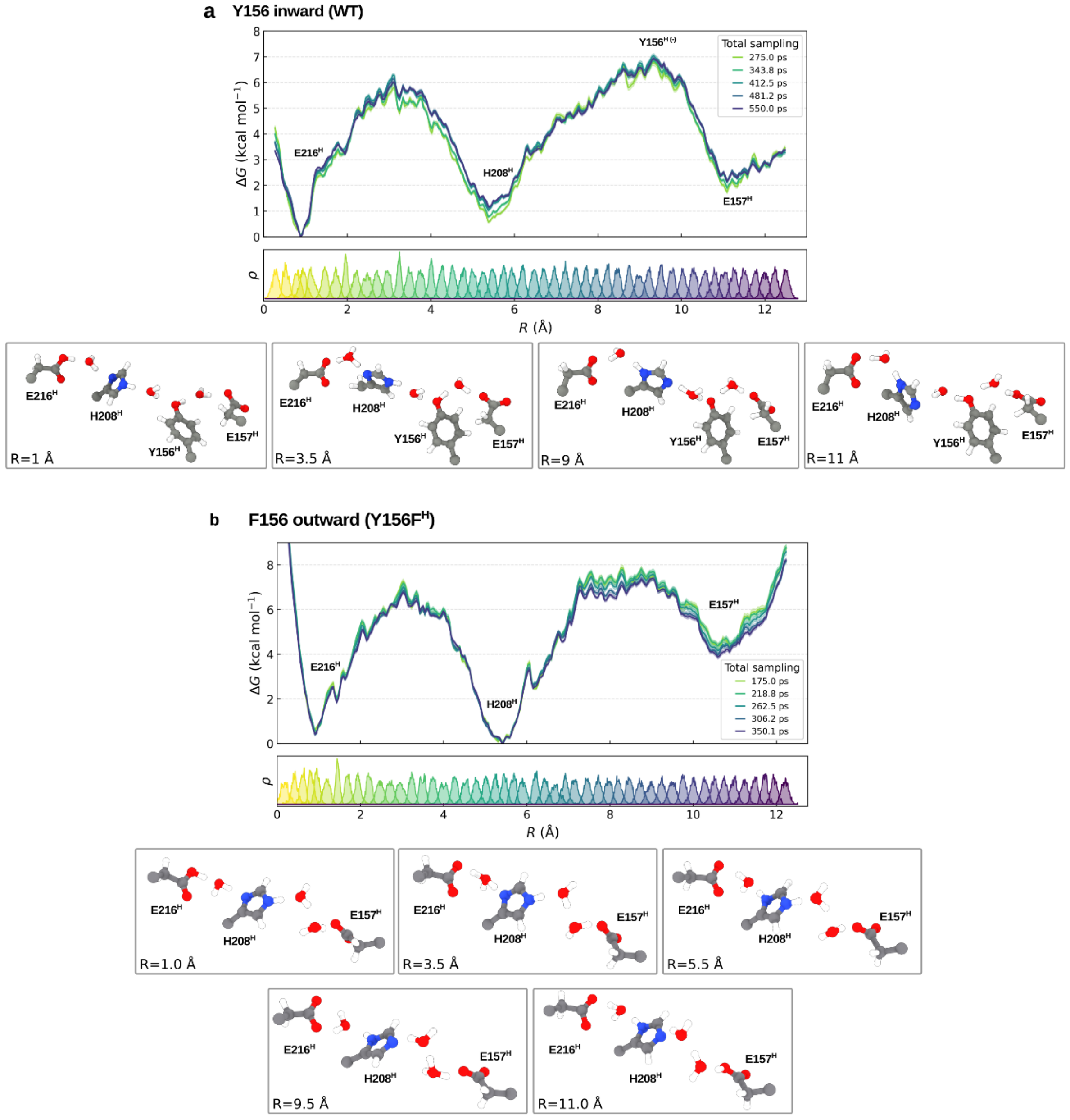


**Supplementary Fig. 5 | Convergence of QM/MM free energy simulations, and structure of intermediates along the proton transfer process.** Free energy profile (*top*) and corresponding histograms of sampled phase space along the reaction coordinate. The reaction coordinate is described by the modified centre of excess charge (mCEC, *middle*), with the principal vector defined from $Glu216^{H}$ to $Glu157^{H}$. The free energies were calculated for **a**, parent Complex I (WT) with *inward*-facing $Tyr156^{H}$ and **b**, the $Y156F^{H}$ variant with *outward*-facing $Phe156^{H}$. Convergence of the free energy profiles is shown by increasing sampling time (colour coded). *Bottom:* representative structures of key intermediates along the QM/MM proton transfer reaction with the reaction coordinate ($R$) labelled in each panel.

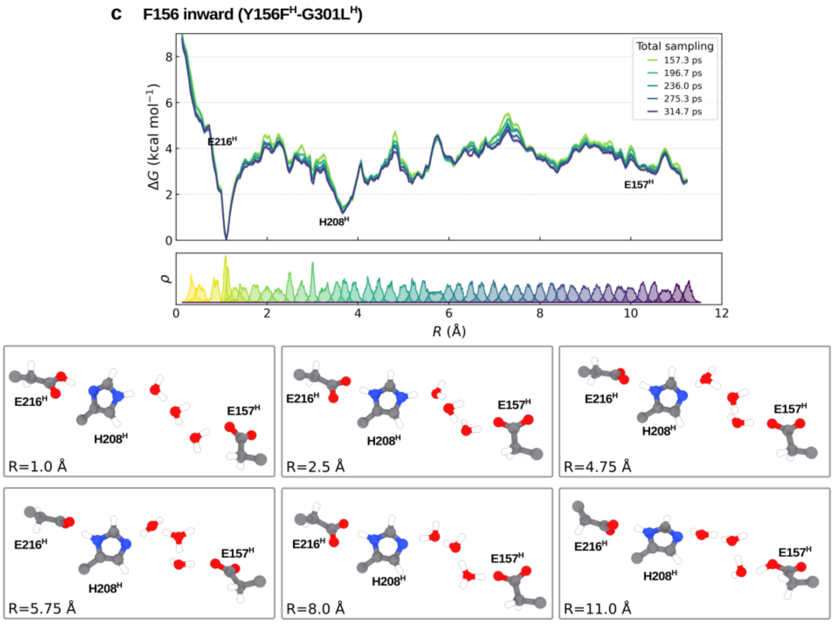


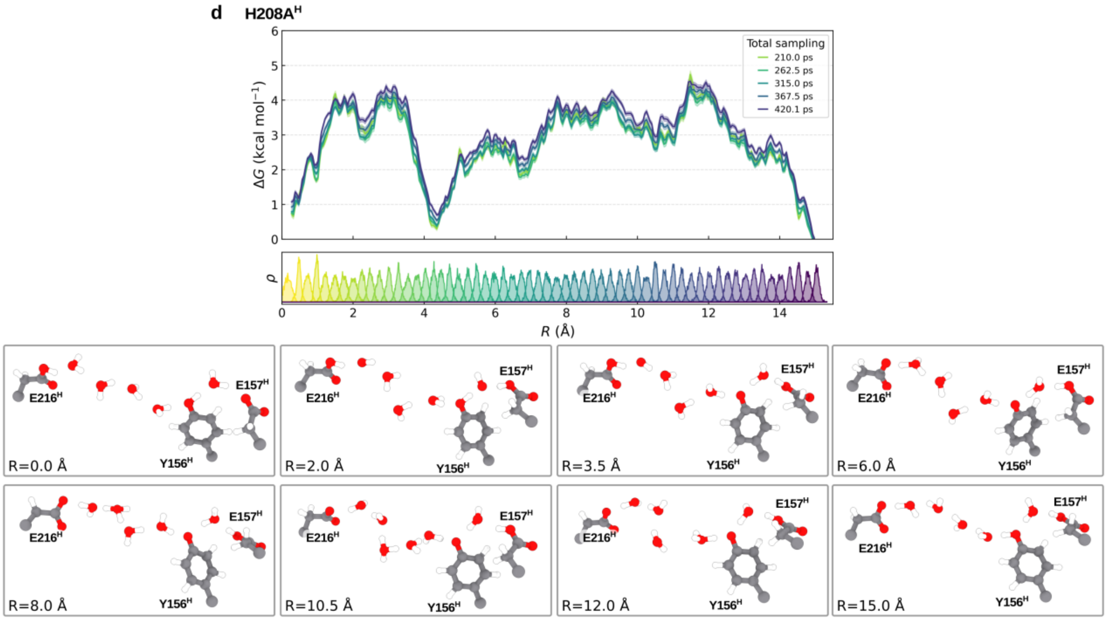


**Supplementary Fig. 5 (*contd.*) | Convergence of QM/MM free energy simulations, and structure of intermediates along the proton transfer process.** Free energy profiles for **c,** the double Y156F$^{H}$-G301L$^{H}$ variant with *inward*-facing Phe156$^{H}$, and **d,** H208A$^{H}$ variant with *inward*-facing Tyr156$^{H}$.

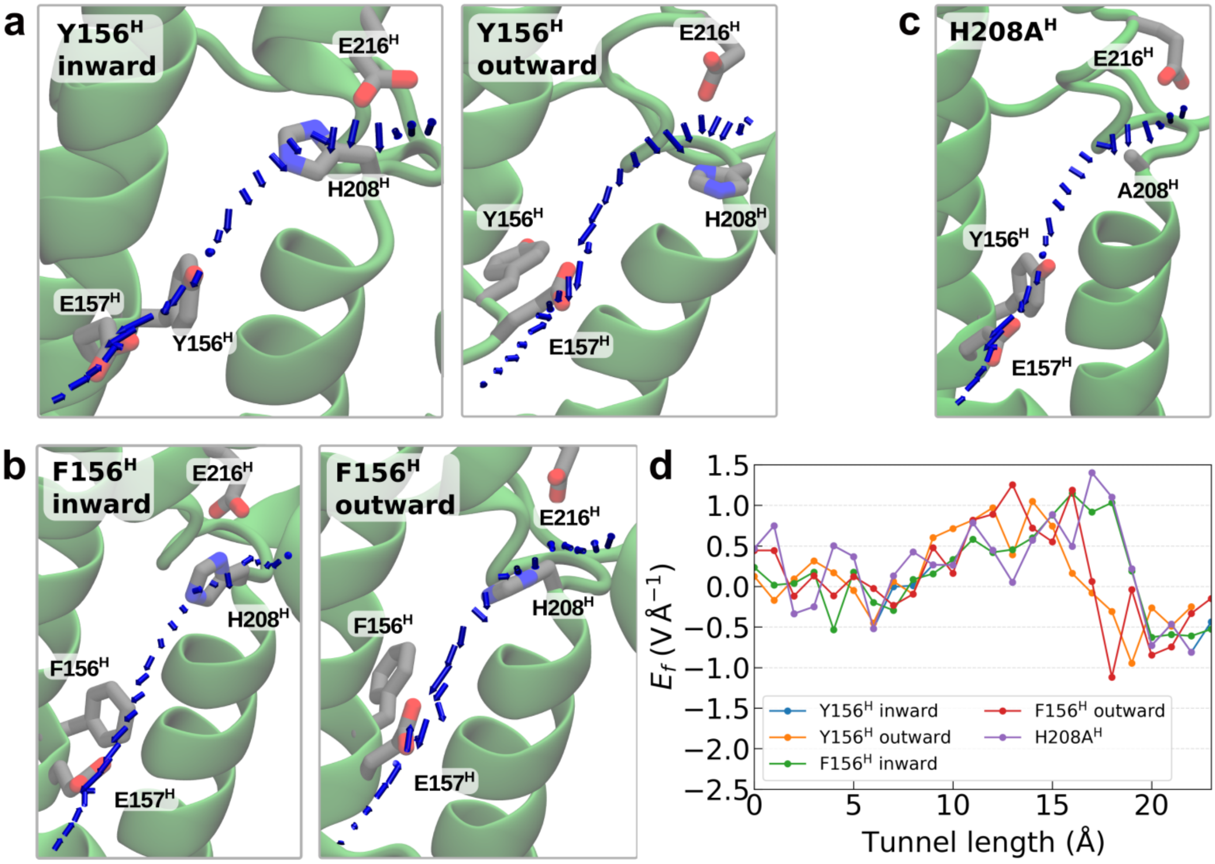


**Supplementary Fig. 6 | Electric field effects along the proton pathway in the WT and Complex I variants.** **a–c,** Representative structures of the proton pathway in different variants and conformational states of Tyr156$^{H}$. Electric field vectors in **a,** for the *inward* and *outward* conformations of Tyr156$^{H}$; **b,** in the *inward* and *outward* conformations of Phe156$^{H}$; and **c,** in the H208A$^{H}$ variant. Blue vectors indicate the local electric field, with direction and vector length indicating relative magnitude. Key residues lining the pathway are shown as sticks. **d,** Quantification of the electric field ($E_f$) projected onto the proton pathway for the NuoH variants and in different conformations of Tyr156$^{H}$. The pathway was defined by tunnel coordinates calculated using CAVER[1].

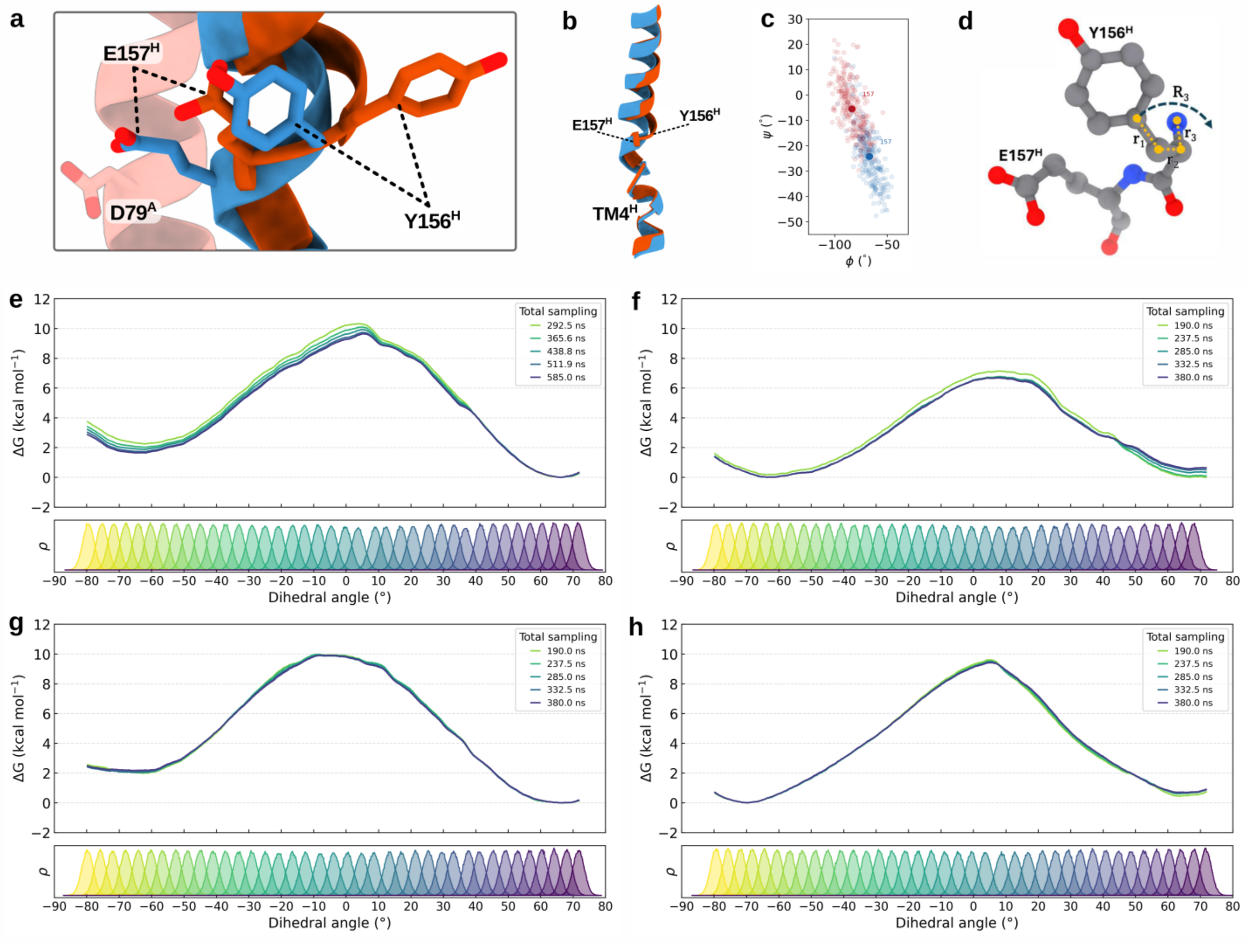


**Supplementary Fig. 7 | Free energy profile of the *inward* → *outward* flip of Tyr156$^{H}$. a,** Structural snapshot showing *inward* and *outward* conformation of Tyr156$^{H}$ from unbiased MD simulations (S1, S3) **b,** Superposition of active TM4$^{H}$ (*blue*) and resting (orange) conformations. **c**, Ramachandran plot of Tyr156$^{H}$ $\varphi$ and $\psi$ dihedral angles extracted from unbiased MD simulations, showing distinct populations of the *inward* and *outward* states. **d**, Reaction coordinate definition used to sample the Tyr156$^{H}$ flip, determined by the dihedral angle formed by C–Cα–Cβ–Cγ atoms. **e–h**, Convergence of free energy profiles for Tyr156$^{H}$ flip for **e**, the parent Complex I (WT); **f**, Y156F$^{H}$; **g**, G301L$^{H}$; and **h**, Y156F$^{H}$/G301L$^{H}$. The free energy profiles at different sampling times are indicated by colour. The homogenous sampling of the reaction phase space is indicated by the histogram (*bottom*).

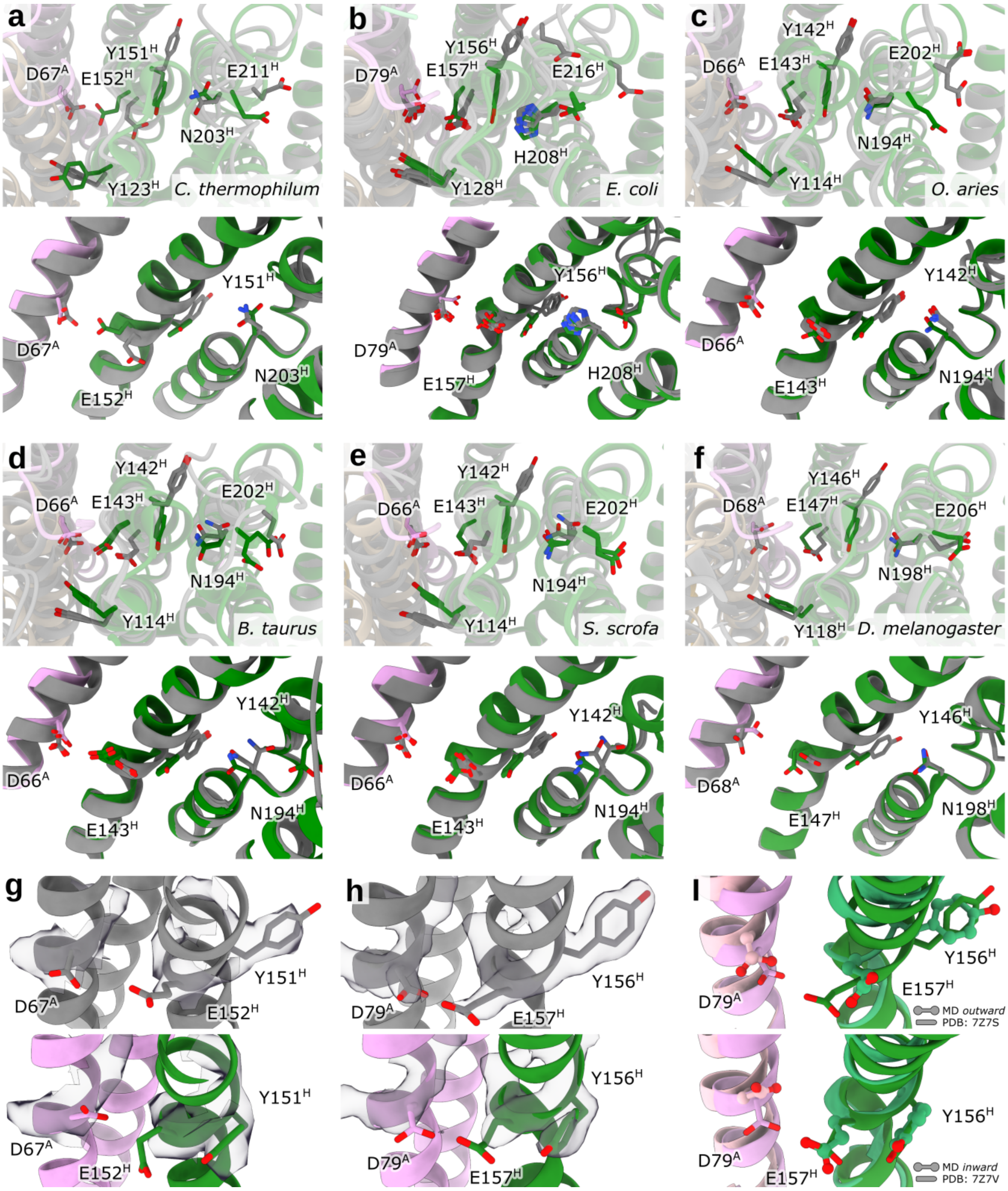


**Supplementary Fig. 8 | Comparison of the *inward*/*outward* conformation of Tyr156$^{H}$ in cryo-EM structures of Complex I. a-f,** Comparison of the E-channel with an α-helix (*coloured*) and π-bulge (in *grey*) at TM3$^{J}$. The structures are from the organisms **a,** *Chaetomium thermophilum* (PDB ID: 7ZMG, 7ZMB); **b,** *Escherichia coli* (PDB ID: 7Z7S, 7Z7V, 7P63, 7P64, 9TAK); **c,** *Ovis aries* (PDB ID: 6ZKC, 6ZKD, 6ZKE, 6ZKF); **d**, *Bos taurus* (PDB ID: 7QSK, 7QSL, 7QSM, 7QSN); **e,** *Sus scrofa* (PDB ID: 7V2H, 7V2K, 7V2C, 7V2D), and **f**, *Drosophila melanogaster* (PDB ID: 8B9Z, 8BA0). **g,h,** Closeup of densities for the E-channel residues of **g,** *D. melanogaster* and **h,** *E. coli* Complex I. **i,** Comparison of *inward* (*top*) and *outward* (*bottom*) conformations from MD simulations with cryo-EM structures in the α- (*top*) and π- (*bottom*) conformations of TM3$^{J}$.

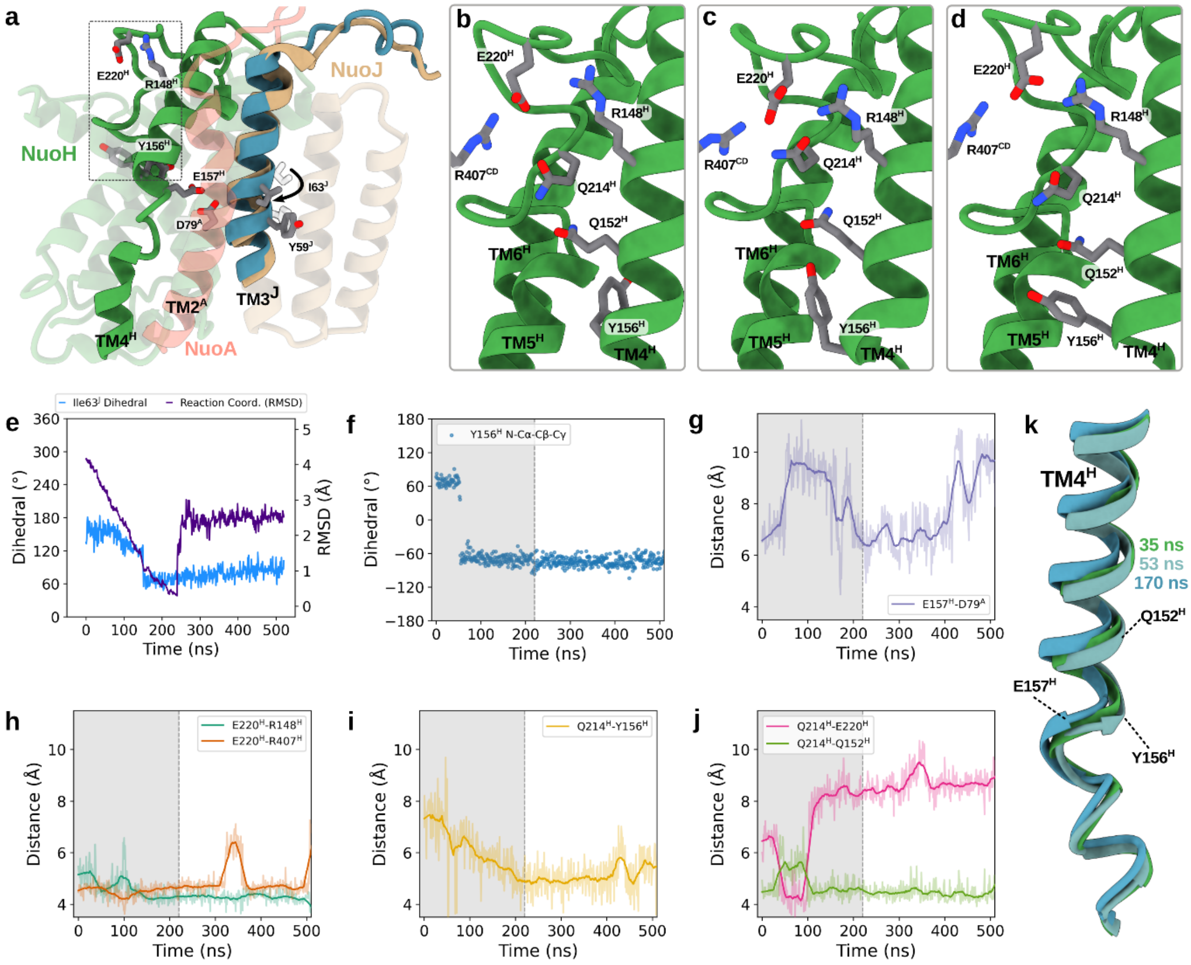


**Supplementary Fig. 9 | Structural re-arrangements from TMD simulation. a,** Overview of key residues, helices, and loops involved in the structural rearrangement during the TMD simulation. The bias was applied on $TM3^J$ for 220 ns, transforming it from an α-helix to the π-bulge. Afterwards, unrestrained MD simulation was performed for *ca.* 380 ns (see *Methods*). **b-d,** Snapshots from TMD simulation at **b,** 35 ns; **c,** 53 ns, and **d,** 170 ns showing the coupled motion of the $Tyr156^H$, $Gln152^H$, $Gln214^H$, $Glu220^H$, $Arg148^H$, and $Arg407^{CD}$. **e,** The dihedral angle (*blue*) between $Phe50^H$ and $Ile63^H$ (Cβ-Cα-Cα-Cβ), and the RMSD between the biased and reference structures during the TMD simulation. **f,** Dihedral angle of $Tyr156^H$ (N-Cα-Cβ-Cγ) describing the *inward* to *outward* flip during the TMD and the subsequent MD simulations. **g-j,** Distance between key residues during the TMD and subsequent MD simulation. The restrained portion of the MD simulation is marked by a grey area. **k,** Snapshots of $TM4^H$ at different timestamps of the TMD simulation.

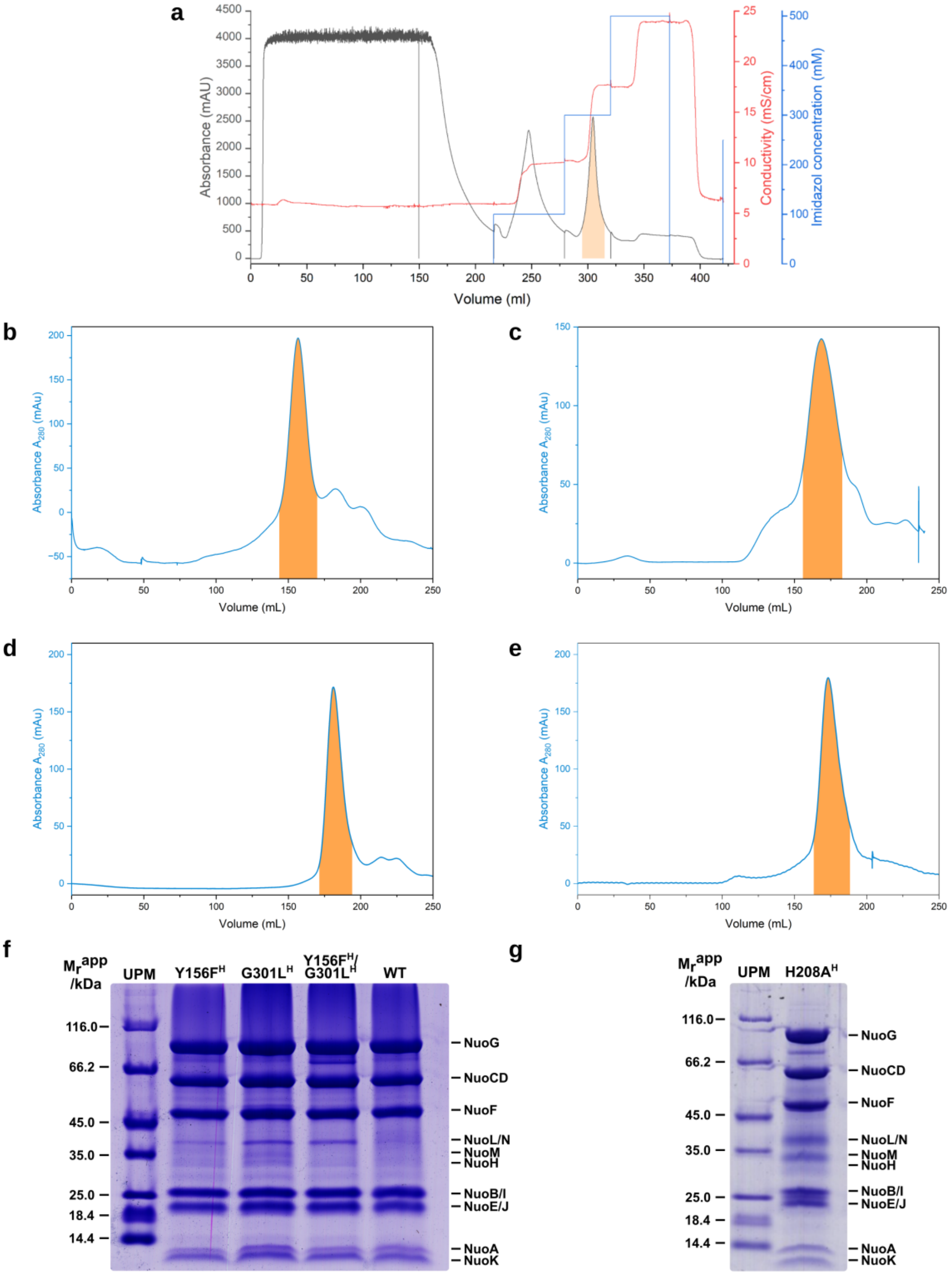


**Supplementary Fig. 10 | Chromatographic profiles for protein purification and SDS-PAGE of all preparations. a,** Representative IMAC chromatogram showing absorbance at 280 nm (*black*), conductivity (*red*), and imidazole concentration (*blue*). The shaded peak indicates the elution fraction collected for further purification. Similar elution profiles were observed for all variants. **b-e,** SEC profiles for the preparations of **b,** Y156F$^{H}$; **c,** G301L$^{H}$; **d,** Y156F$^{H}$/G301L$^{H}$ and **e,** H208A$^{H}$ variants. Absorbance at 280 nm is shown. The shaded peak indicates the collected elution fraction. **f,** SDS-PAGE of the preparations of the Y156F$^{H}$, G301L$^{H}$, Y156F$^{H}$/G301L$^{H}$ variants and parent complex (WT), confirming the presence of all Nuo subunits. **g,** SDS–PAGE analysis of the preparation of the H208A$^{H}$ variant.

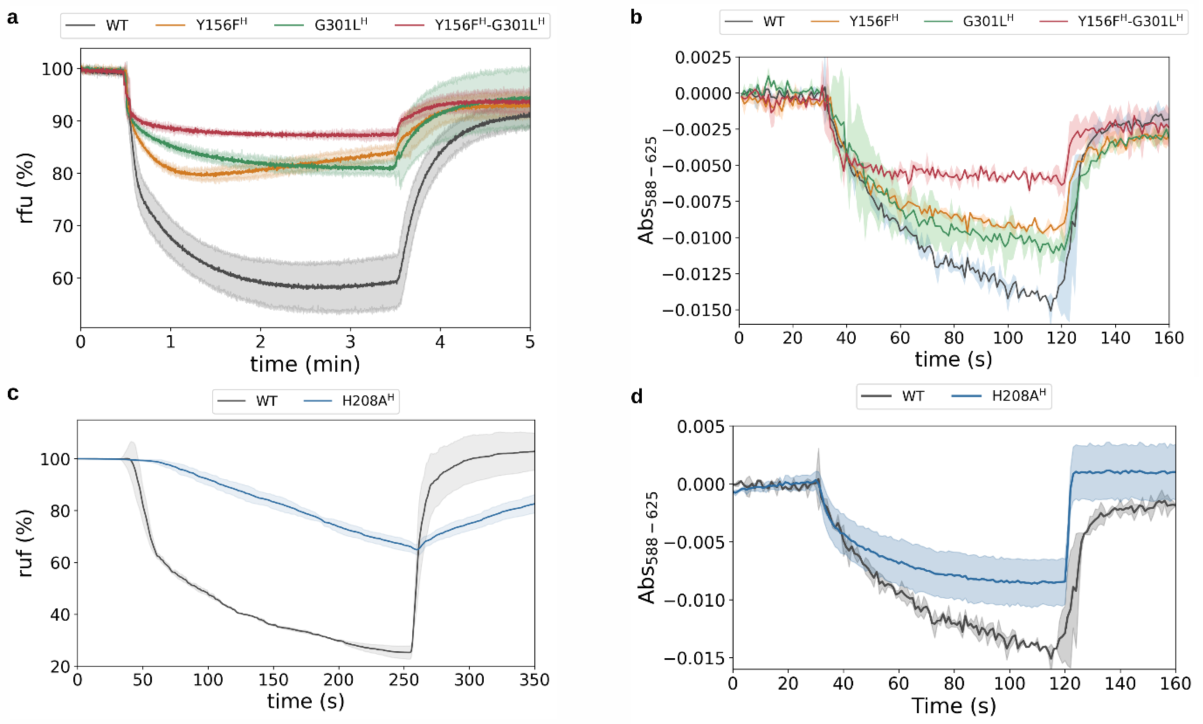


**Supplementary Fig. 11 | Proton translocation and membrane potential formation in the NuoH variants. a,** ACMA fluorescence quenching assay monitoring ΔpH formation by proteoliposomes of parent Complex I (WT), Y156F$^{H}$, G301L$^{H}$, and Y156F$^{H}$/G301L$^{H}$ variants. Traces are normalized to the initial fluorescence signal. **b,** Oxonol VI absorbance change (ΔAbs$_{588-625}$) reporting membrane potential (ΔΨ) generation for the same set of variants. **c,** ACMA fluorescence quenching by the H208A$^{H}$ variant (*red*) and parent Complex I (WT, *black*). **d,** Generation of a membrane potential by the H208A$^{H}$ variant (*orange*) and the parent Complex I (WT, *blue*).

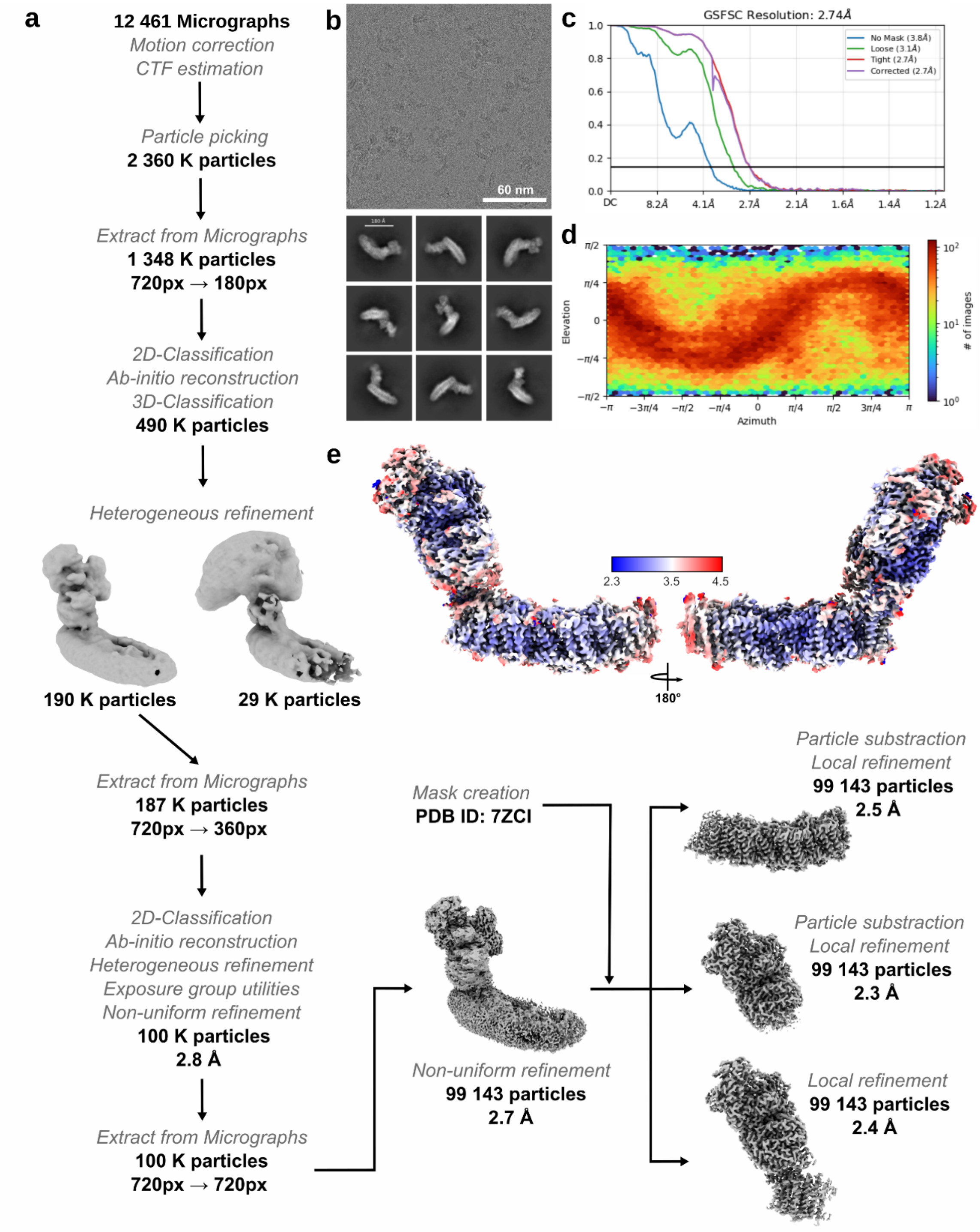


**Supplementary Fig. 12 | Cryo-EM data analysis and validation of the 2.7 Å map of the Y157F$^{H}$ variant. a,** Schematic overview of the cryo-EM data processing workflow. A total of 12,461 micrographs were used for particle picking and iterative refinement, yielding a final particle stack of 99,143 particles. **b,** Representative motion-corrected micrograph and selected 2D class averages. **c,** Global FSC curve of the final non-uniform refinement with a corrected resolution of 2.74 Å. **d,** Angular distribution plot showing the orientation distribution. **e**, Local resolution map of the non-uniform refined model, indicating highest resolution (~2.3 Å) in the core of the membrane arm and lower resolution in peripheral regions. Local refinements of membrane and hydrophilic arms yielded maps at 2.5 Å, 2.3 Å, and 2.4 Å, respectively.

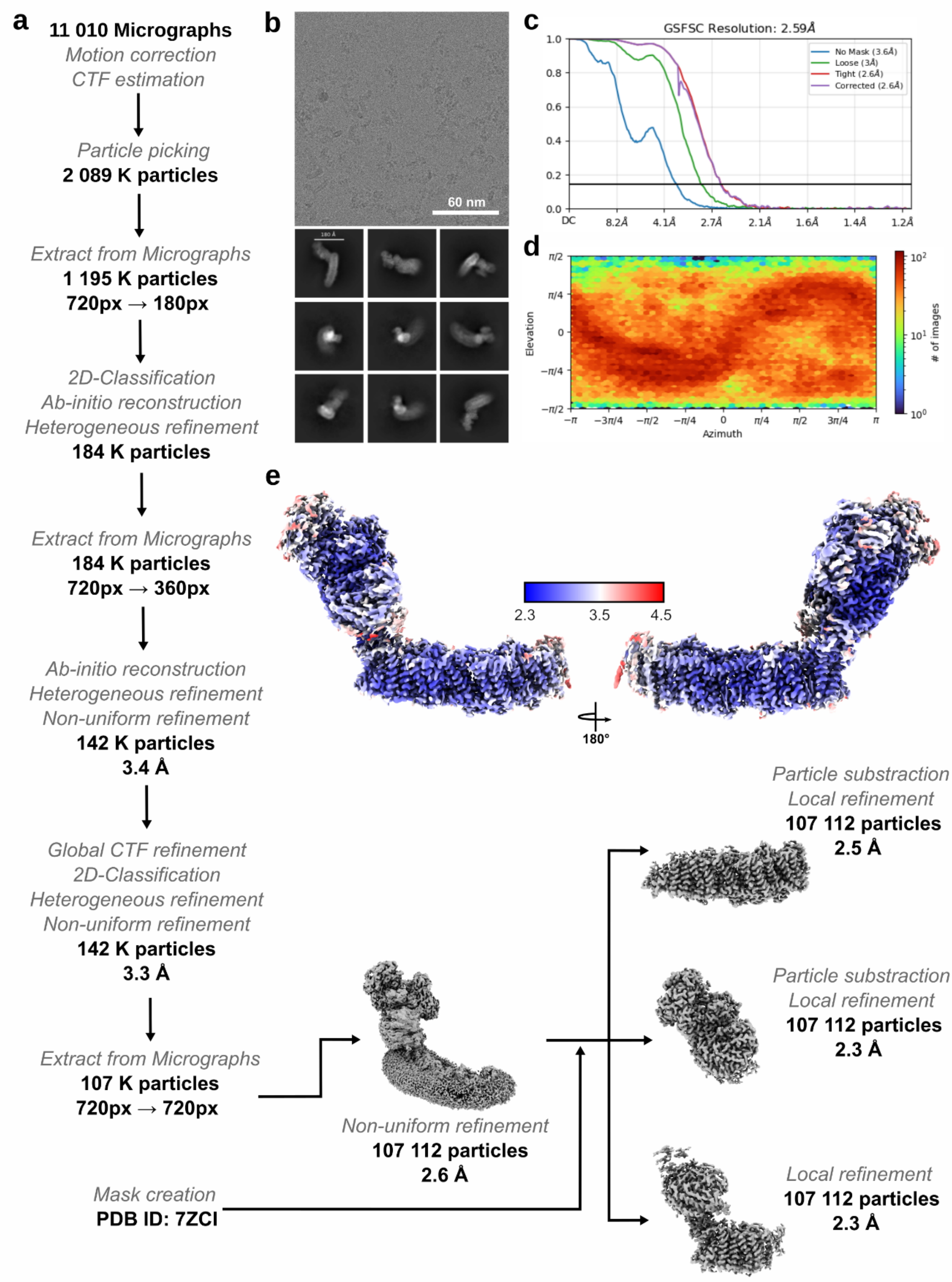


**Supplementary Fig. 13 | Cryo-EM data analysis and validation of the 2.6 Å map of the Y157F$^{H}$/G301L$^{H}$ variant. a**, Workflow of cryo-EM data processing. A total of 11,010 micrographs were collected and processed, resulting in 107,112 particles used for final refinement. **b**, Representative micrograph and selected 2D class averages. **c**, Gold-standard FSC curve of the final non-uniform refinement, indicating a corrected global resolution of 2.59 Å. **d**, Angular distribution plot showing even particle orientation. **e**, Local resolution representation of the consensus map. Local refinements following particle subtraction yielded maps of the membrane and hydrophilic domains at 2.5 Å, 2.3 Å, and 2.3 Å, respectively.

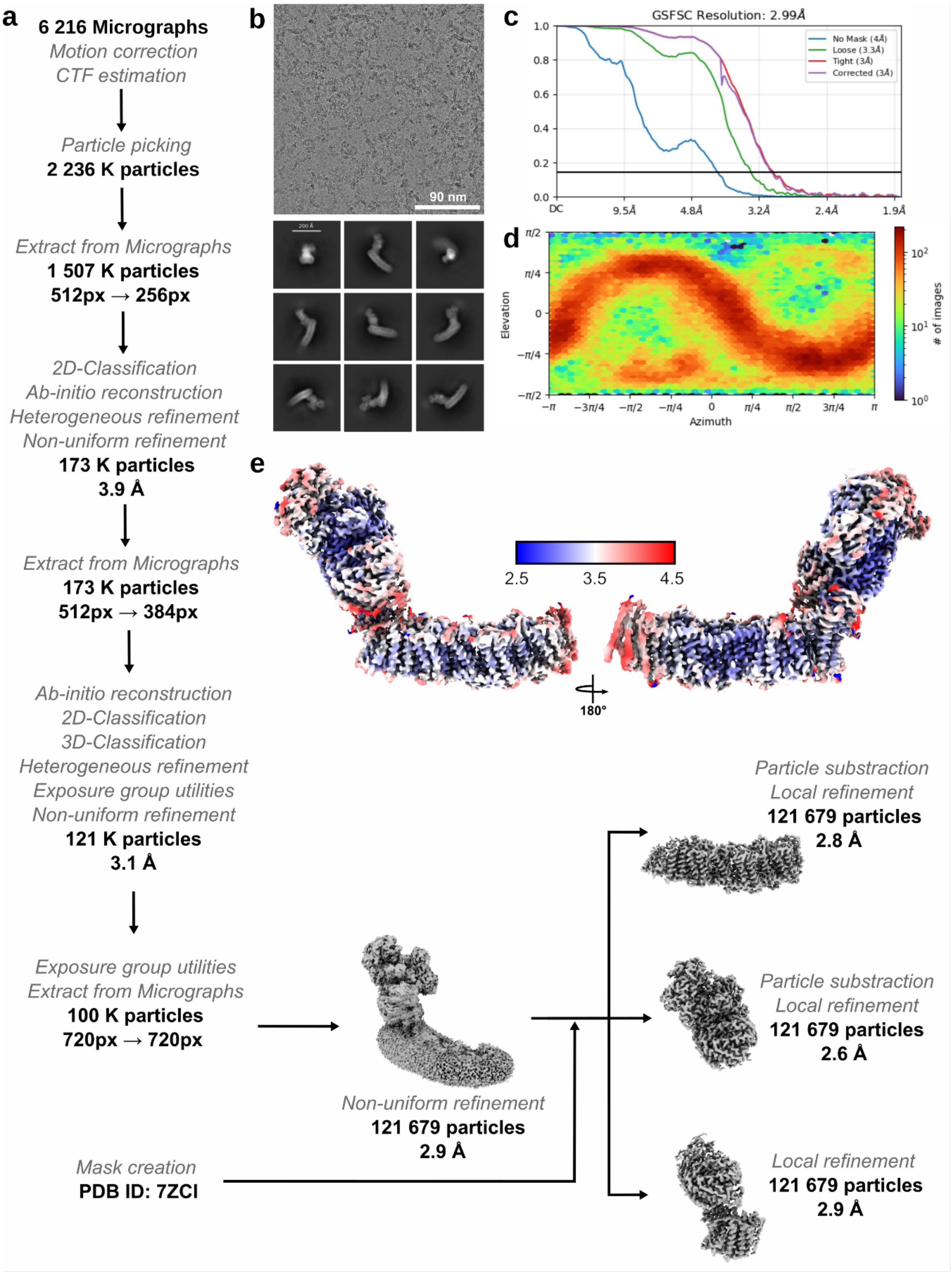


**Supplementary Fig. 14 | Cryo-EM data analysis and validation of the 2.9 Å map of the G301L$^{H}$ variant. a,** Cryo-EM data processing workflow. A total of 6,216 micrographs were collected and processed, yielding 121,679 particles for final refinement. **b**, Representative micrograph and selected 2D class averages. **c**, Gold-standard FSC curve of the non-uniform refinement, with a corrected resolution of 2.99 Å. **d**, Angular distribution plot showing particle orientations. **e**, Local resolution distribution of the final map. Local refinements after particle subtraction resulted in membrane and hydrophilic domain maps at 2.8 Å, 2.6 Å, and 2.9 Å, respectively.

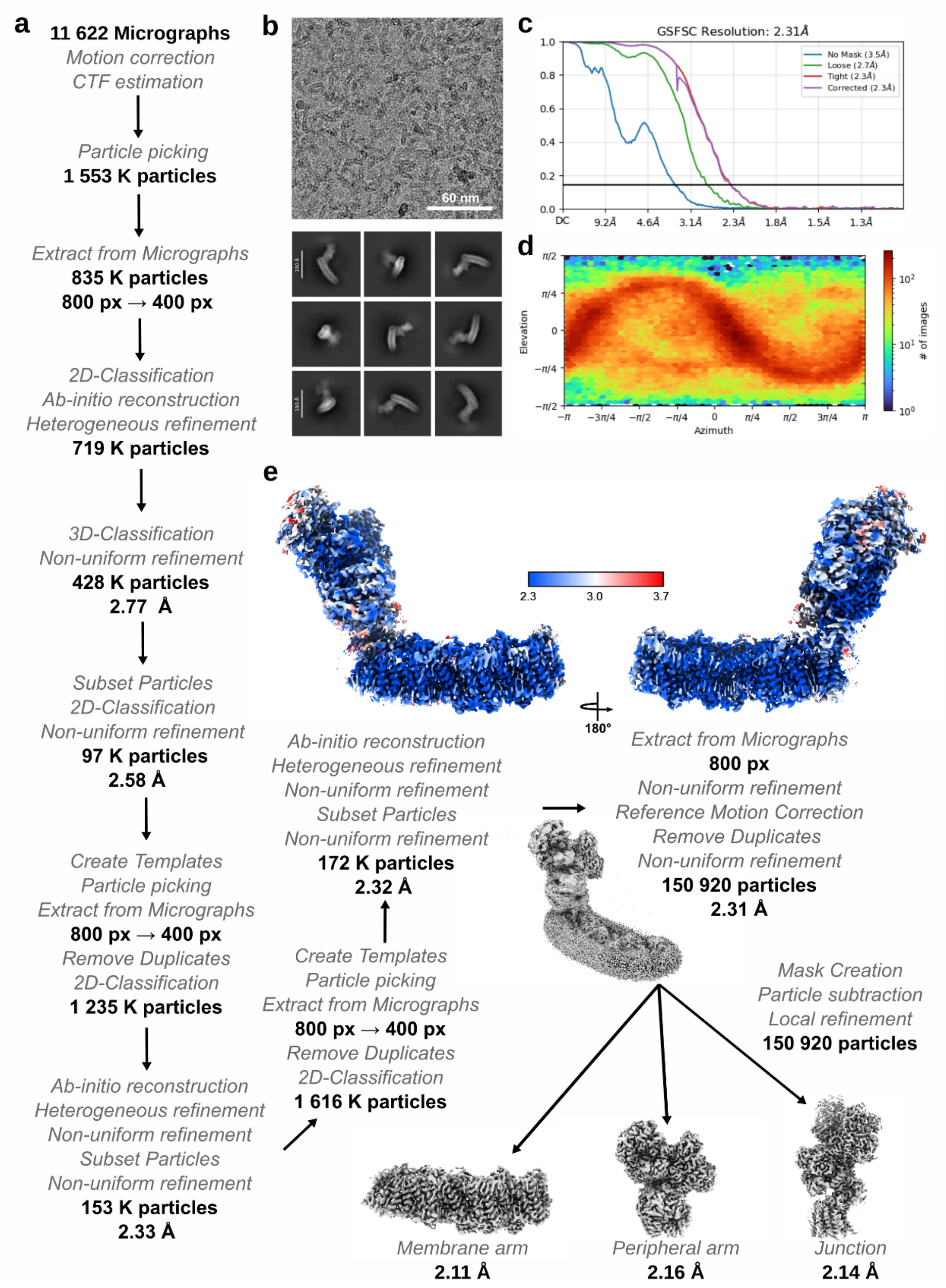


**Supplementary Fig. 15 | Cryo-EM data analysis and validation of the 2.3 Å map of the H208A$^{H}$ variant.** **a,** Cryo-EM data processing workflow. A total of 11,622 micrographs were motion-corrected and CTF-estimated, yielding ~1.55 million initially picked particles. After iterative rounds of extraction, 2D classification, *ab initio* reconstruction, heterogeneous and non-uniform refinement, particle subsetting, and template-based repicking, 150,920 particles were retained for the final non-uniform refinement. **b**, Representative micrograph and selected 2D class averages showing well-defined particle features. **c**, Gold-standard FSC curves from non-uniform refinement, with a corrected global resolution of 2.31 Å (FSC=0.143 criterion). **d**, Angular distribution of particles used in the final reconstruction, demonstrating broad orientation coverage. **e**, Final cryo-EM density map coloured by local resolution. Focused refinements following particle subtraction yielded maps of the membrane arm (2.11 Å), peripheral arm (2.16 Å), and junction region (2.14 Å).

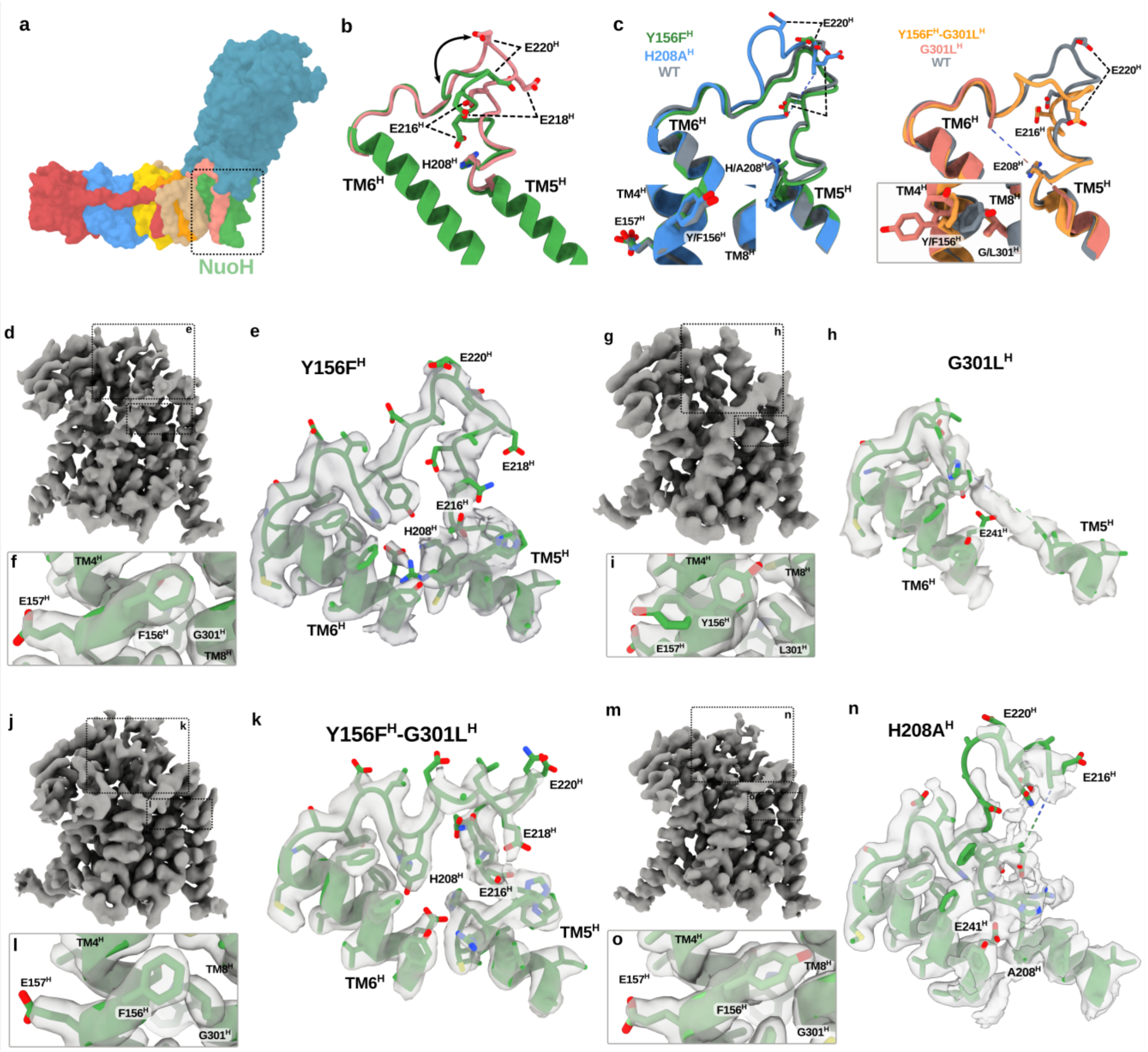


**Supplementary Fig. 16 | Example densities based on the 2.2-2.9 Å maps of the variants. a,** Surface representation of Complex I showing the NuoH subunit (green), comprising the TM5-6$^{H}$ loop shown in panels below. **b,** Proposed conformational rearrangement within NuoH involving TM5$^{H}$, TM6$^{H}$, and conserved glutamate residues (Glu216$^{H}$, Glu218$^{H}$, Glu220$^{H}$) and His208$^{H}$ based on structures obtained. **c,** Superposition of the TM5-6$^{H}$ loop from the Y156F$^{H}$ (green) and H208A$^{H}$ (teal) variants as compared to the Complex I resting-state structure (PDB ID: 9TAK, blue), highlighting differences in loop conformation and sidechain positions. **d–f,** Cryo-EM map (transparent surface) and atomic model of the TM5-6$^{H}$ region for the Y156F$^{H}$ variant. **d,** Overview of the local cryo-EM map for NuoH. **e,** Fit of the atomic model of the TM5-6$^{H}$ loop. **f,** Close-up showing the orientation of F156$^{H}$. **g–i,** Cryo-EM map and atomic model of the TM5-6$^{H}$ region for the G301L$^{H}$, **j-l,** the Y156F$^{H}$/G301L$^{H}$ and **m–o,** the H208A$^{H}$ variant. **i,** Alternative conformation of Tyr156$^{H}$.

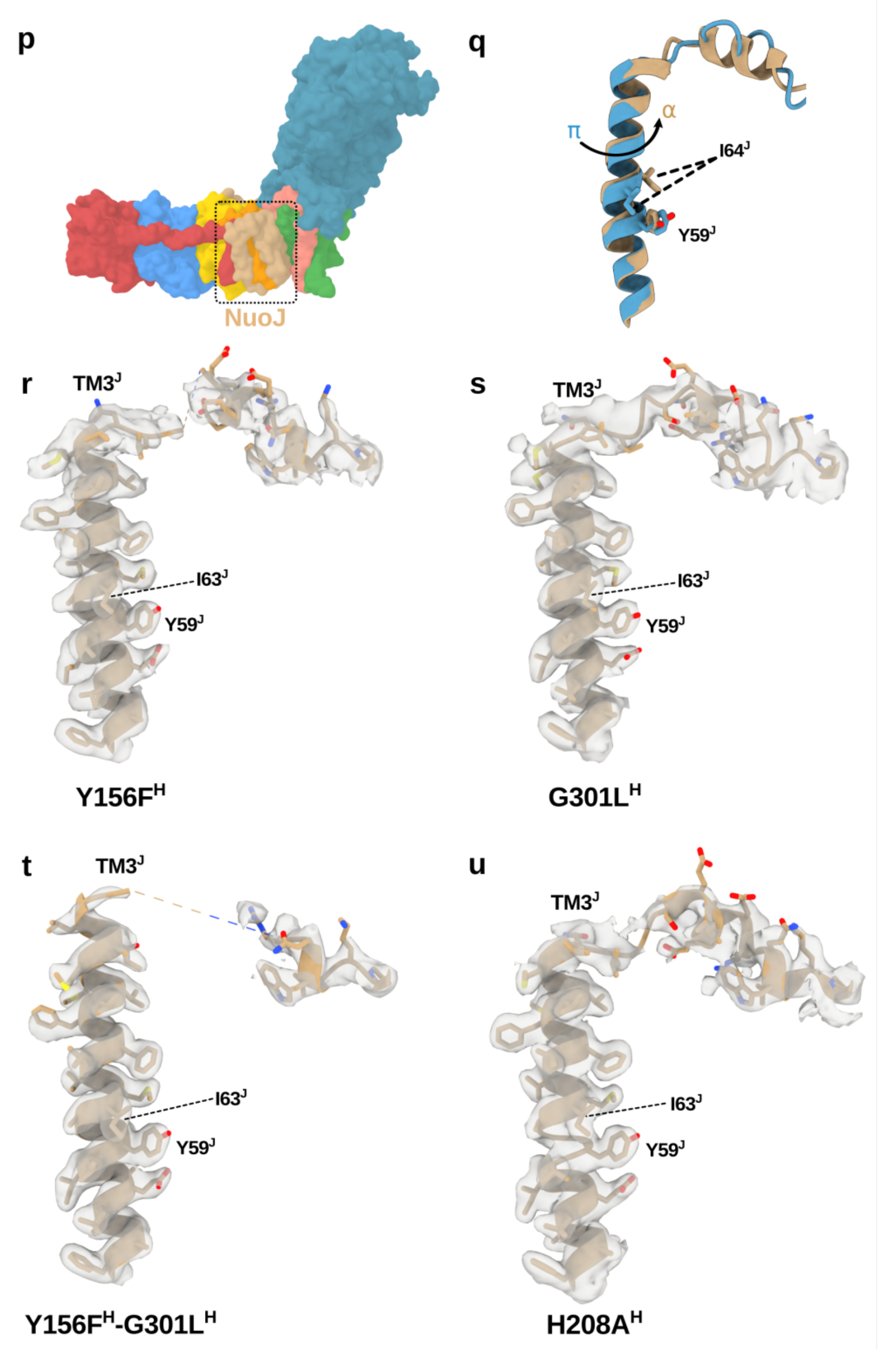


**Supplementary Fig. 16 (contd.) | Example densities based on the 2.2-2.9 Å maps of the variants. p**, Surface representation of Complex I highlighting NuoJ (orange) that comprises helix $TM3^J$ shown in the other panels. **q**, The $\pi \rightarrow \alpha$ transition of $TM3^J$ comparing the $\alpha$-helical conformation (blue) and the $\pi$-bulge (orange), and reposition of $Tyr59^J$ and $Ile64^J$. **r–u,** Cryo-EM maps (transparent mesh) and fitted atomic models of $TM3^J$ and the connecting loop to the adjacent helix for the **r**, $Y156F^H$**, s**, $G301L^H$, **t**, $Y156F^H/G301L^H$**,** and **u**, $H208A^H$ variants.

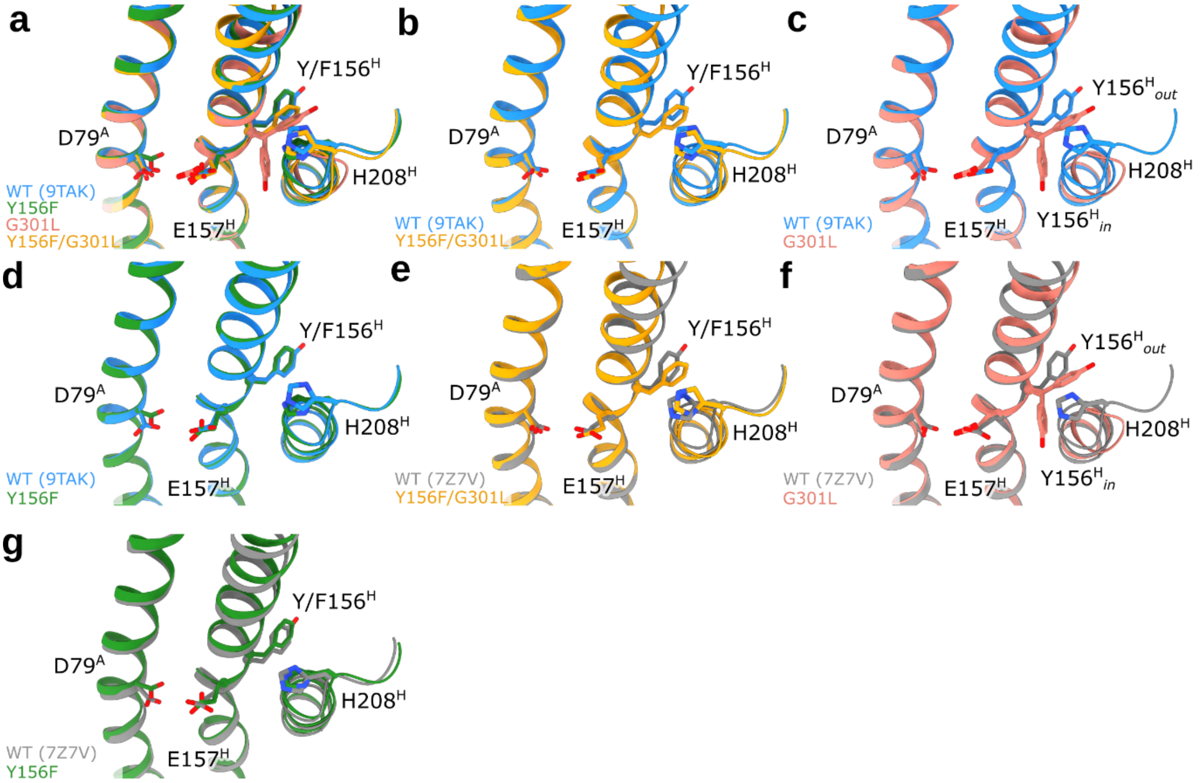


**Supplementary Fig. 17 | Structure of different conformational states of Tyr156$^H$ and Glu157$^H$ in various NuoH variants. a–d,** Superposition of the TM4$^H$–TM6$^H$ region from cryo-EM structures comparing parent Complex I (WT; PDB: 9TAK[33], *blue*) with the Y156F$^H$ (*green*), G301L$^H$ (*red*), and Y156F$^H$/G301L$^H$ (*orange*) variants. **e–g,** Comparison of the variant structures with the WT resting-state structure (PDB ID: 7Z7V[12], *grey*).

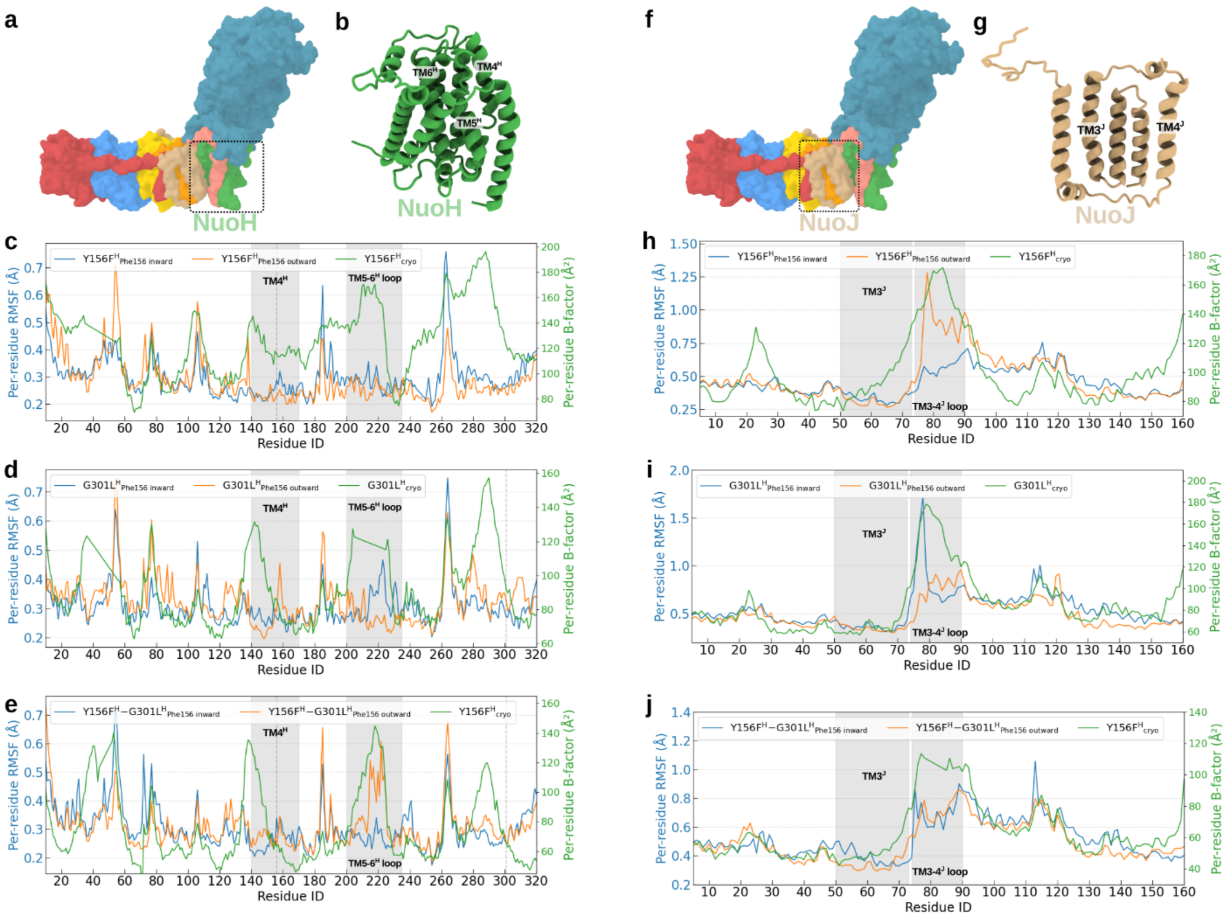


**Supplementary Fig. 18 | Comparison of RMSF from MD simulations with beta factors from cryo-EM data.** **a,** Surface representation of *E. coli* Complex I highlighting NuoH (green). **b,** Cartoon representation of NuoH, indicating the positions of TM4$^{H}$, TM5$^{H}$, TM6$^{H}$, and the TM5–6$^{H}$ loop. **c–e,** Root Mean Square Fluctuation (RMSF, computed on Cα positions) from 500 ns MD simulations of the *inward* (blue) or *outward* (orange) conformation of **c**, Y156F$^{H}$, **d**, G301L$^{H}$, and **e**, Y156F$^{H}$/G301L$^{H}$ variants. Shaded regions highlight TM4$^{H}$ and the TM5–6$^{H}$ loop and the dashed line the mutation site. **f,** Surface representation of *E. coli* Complex I highlighting the NuoJ subunit (*beige*). **g,** Cartoon representation of NuoJ, showing TM3$^{J}$ and TM4$^{J}$. **h–j,** RMSF profiles for NuoJ from MD simulations of the **h**, Y156F$^{H}$, **i**, G301L$^{H}$, and **j**, Y156F$^{H}$/G301L$^{H}$ variants, shown together with cryo-EM per-residue B-factors (*green*).

**Supplementary Table 1 | List of performed MD simulations and modelled states.** A quinol anion ($QH^-$) was modelled in the Q site 2.

| Simulation | Variant | Substrate | Conformational State | Length (ns) |
|---|---|---|---|---|
| S1 | WT | $QH^-$ | *inward* $Tyr156^H$ | 500 |
| S2 | WT | $QH^-$ | *inward* $Tyr156^H$ | 500 |
| S3 | WT | $QH^-$ | *outward* $Tyr156^H$ | 500 |
| S4 | WT | $QH^-$ | *outward* $Tyr156^H$ | 500 |
| S5 | $Y156F^H$ | $QH^-$ | *inward* $Phe156^H$ | 500 |
| S6 | $Y156F^H$ | $QH^-$ | *inward* $Phe156^H$ | 500 |
| S7 | $Y156F^H$ | $QH^-$ | *outward* $Phe156^H$ | 500 |
| S8 | $Y156F^H$ | $QH^-$ | *outward* $Phe156^H$ | 500 |
| S9 | $G301L^H$ | $QH^-$ | *outward* $Tyr156^H$ | 500 |
| S10 | $G301L^H$ | $QH^-$ | *inward* $Tyr156^H$ | 500 |
| S11 | $G301L^H$ | $QH^-$ | *outward* $Tyr156^H$ | 500 |
| S12 | $G301L^H$ | $QH^-$ | *outward* $Tyr156^H$ | 500 |
| S13 | $Y156F^H$-$G301L^H$ | $QH^-$ | *inward* $Phe156^H$ | 500 |
| S14 | $Y156F^H$-$G301L^H$ | $QH^-$ | *inward* $Phe156^H$ | 500 |
| S15 | $Y156F^H$-$G301L^H$ | $QH^-$ | *outward* $Phe156^H$ | 500 |
| S16 | $Y156F^H$-$G301L^H$ | $QH^-$ | *outward* $Phe156^H$ | 500 |
| S17 | $H208A^H$ | $QH^-$ | *inward* $Tyr156^H$ | 500 |
| S18 | $H208A^H$ | $QH^-$ | *inward* $Tyr156^H$ | 500 |
| S19 | WT | $QH^-$ | *inward* $Tyr156^H$, Glu216 (-) | 500 |
| S20 | WT | $QH^-$ | TMD, *inward* → *outward* $Tyr156^H$ | 520 |
| S21 | WT | $QH_2$ | *inward* $Tyr156^H$ | 125 |
| **Total:** | | | | **10.14 µs** |

**Supplementary Table 2 | List of performed QM/MM free energy simulations.**

| **Simula-tion** | **Starting structure** | **Variant** | **QM region** | **Atoms *N*** | **Sampling** |
|---|---|---|---|---|---|
| Q1 | S1 | WT | Glu126$^{H}$, His208$^{H}$, Tyr156$^{H}$, Glu157$^{H}$, Val206$^{H}$, Val127$^{H}$, Thr153$^{H}$, Gln152$^{H}$, Tyr225$^{H}$, 12 $H_2O$ | 138 | 50 x 6 ps |
| Q2 | S5 | Y156F$^{H}$ | Glu126$^{H}$, His208$^{H}$, Glu157$^{H}$, Val206$^{H}$, Val127$^{H}$, Thr153$^{H}$, Gln152$^{H}$, Tyr225$^{H}$, Ala149$^{H}$ 11 $H_2O$ | 121 | 50 x 4 ps |
| Q3 | S13 | Y156F$^{H}$/ G301L$^{H}$ | Glu126$^{H}$, His208$^{H}$, Phe156$^{H}$, Glu157$^{H}$, Val206$^{H}$, Val127$^{H}$, Thr153$^{H}$, Gln152$^{H}$, Tyr225$^{H}$, Ala149$^{H}$, 18 $H_2O$ | 156 | 46 x 6 ps |
| Q4 | S17 | H208A$^{H}$ | Glu126$^{H}$, Ala208$^{H}$, Tyr156$^{H}$, Glu157$^{H}$, Val206$^{H}$, Val127$^{H}$, Thr153$^{H}$, Gln152$^{H}$, Tyr225$^{H}$, Ala149$^{H}$, 19 $H_2O$ | 153 | 59 x 4 ps |
| **Total** | | | | | 1012 ps |

**Supplementary Table 3 | List of performed classical free energy simulations.**

| **Simu-lation** | **Starting structure** | **Variant** | **Sampling range (Y/F156 $^{H}$ dihedral)** | **Atoms *N*** | **Sampling** |
|---|---|---|---|---|---|
| C1 | S1 | WT | -80° - 72° | 853 241 | 39 x 15 ns |
| C2 | S5 | Y156F$^{H}$ | -80° - 72° | 853 240 | 39 x 10 ns |
| C3 | S9 | G301L$^{H}$ | -80° - 72° | 853 253 | 39 x 10 ns |
| C4 | S13 | Y156F$^{H}$/ G301L$^{H}$ | -80° - 72° | 853 252 | 39 x 10 ns |
| **Total** | | | | | **1.75 µs** |

**Supplementary Table 4 | List of base protonation states used in the MD simulations.** Only non-standard protonation states are listed. His(δ) – $N_δ$ protonated (neutral) histidine, His(ε) – $N_ε$ protonated (neutral) histidine, (ε/δ) – ε/δ protonated (charged) histidine. Unless otherwise stated, the histidine residues were modelled in their $N_δ$ protonated (neutral) form.

| Subunit | Residues |
|---|---|
| NuoA | Asp79, Glu81, Glu102 |
| NuoB | - |
| NuoCD | His78(ε), His87(δ/ε), His101(ε), His123(δ/ε), Glu138, His152(ε), His224(ε), His228(δ/ε), Asp273, His319(ε), His511(ε), His518(ε), Glu538, His568(ε), Asp597, Asp599 |
| NuoE | Glu25, His56(ε), His87(δ/ε) |
| NuoF | NuoF His11(δ), Asp92, Glu93, His110(ε), His172(δ), Asp298, His304(δ), His331(ε), Glu343, His400(δ), His431(ε) |
| NuoG | Glu23, Glu110, His115(ε), Asp118, His125(δ/ε), His178(ε), Glu243, Glu252, His261(ε), Asp266, Glu382, Asp383, Glu570 His576(δ/ε), Glu707, His819(ε) |
| NuoH | His208(ε), His210(ε), Glu218, Glu241 His226(ε) |
| NuoI | Glu124 |
| NuoJ | His27(δ/ε), Glu55, Glu142, His157(ε) |
| NuoK | His84(ε) |
| NuoL | His207(δ/ε), Lys399, Glu587 |
| NuoM | Glu108, Asp135, His241(δ/ε), His322(ε), His348(ε) |
| NuoN | His34(ε), His305(ε), Lys395 |

**Supplementary Table 5 | Cryo-EM data collection, refinement, and validation statistics.**

| **Data collection, processing** | **Dataset 1: Y156F$^{H}$** PDB ID: 28WM, EMBD: 56915, 56916, 56917, 56918, 56913 | **Dataset 2: Y156F/G301L$^{H}$** PDB ID: 28TL, EMBD: 56811, 56812, 56815, 56817, 56818 | **Dataset 3: G301L$^{H}$** PDB ID: 28VI, EMBD: 56880, 56881, 56882, 56883, 56884 | **Dataset 3: H208A$^{H}$** PDB ID: 28NI, EMBD: 56649 56653, 56654, 56655, 56656 |
|---|---|---|---|---|
| Voltage (kV) | 300 | 300 | 300 | 300 |
| Magnification | 215 000 | 215 000 | 130 000 | 215 000 |
| Electron exposure ($e^{-}$/Å$^{2}$) | 40 | 40 | 40 | 40 |
| Pixel size (Å) | 0.572 | 0.572 | 0.96 | 0.572 |
| Defocus range (µm) | -0.5 to -2.3 | -0.5 to -2.0 | -0.5 to -2.0 | -0.5 to 2.2 |
| Defocus step (µm) | -0.2 | -0.2 | -0.2 | 0.1 |
| Symmetry imposed | None (C1) | None (C1) | None (C1) | None (C1) |
| Initial particle images (number) | 2 360 000 | 2 089 000 | 2 236 000 | 1 553 479 |
| Final particle images (number) | 99143 | 107112 | 121 679 | 150 920 |
| FSC threshold | 0.143 | 0.143 | 0.143 | 0.143 |
| Map resolution (Å) | 2.7 | 2.5 | 2.9 | 2.3 |
| **Refinement** | | | | |
| CC (mask) | 0.90 | 0.89 | 0.89 | 0.89 |
| Resolution estimates (Å) | | | | |
| d 99 | | | | |
| Masked | 2.7 | 2.6 | 3.1 | 2.4 |
| Unmasked | 2.4 | 2.3 | 2.8 | 2.2 |
| d FSC model, 0/0.143/0.5 (Å) | | | | |
| Masked | 1.8/1.9/2.8 | 1.7/1.9/2.8 | 1.9/2.0/3.0 | 1.4/1.7/2.4 |
| Unmasked | 1.8/2.0/2.9 | 1.7/2.0/2.8 | 1.9/2.1/3.1 | 1.4/1.8/2.6 |
| Model composition | | | | |
| Protein residues | 4692 | 4707 | 4666 | 4733 |
| Non-hydrogen atoms | 38057 | 38003 | 37502 | 28293 |
| Ligands | 37 | 6740 | 36 | 25 |
| Water | 584 | 24916 | 80 | 777 |
| **B factors (Å$^{2}$)** | | | | |
| **Min/max/mean** | | | | |
| Protein residues | 23.36/205.68/71.71 | 30.00/171.13/73.88 | 41.04/202.18/89.61 | 25.12/152.36/53.41 |
| Ligands | 47.34/163.37/99.86 | 20.00/165.48/68.83 | 61.73/194.42/115.51 | 20.00/140.29/58.48 |
| Water | 30.00/132.33/63.68 | 30.00/99.95/59.77 | 58.31/109.15/81.42 | 10.00/50.82/12.00 |
| **Validation** | | | | |
| MolProbity | 1.34 | 1.57 | 1.62 | 1.61 |
| Root mean square deviations | | | | |
| Bond lengths (Å) | 0.002 | 0.010 | 0.005 | 0.015 |
| Bond angles (°) | 0.498 | 1.081 | 0.564 | 1.693 |
| Clash score | 5.37 | 7.36 | 6.91 | 7.49 |
| Poor rotamers (%) | 0.94 | 0.89 | 1.10 | 0.93 |
| **Ramachandran plot** | | | | |
| Favoured (%) | 97.76 | 97.06 | 96.73 | 96.81 |
| Allowed (%) | 2.24 | 2.89 | 3.27 | 3.13 |
| Disallowed (%) | 0.00 | 0.04 | 0.00 | 0.06 |

**Supplementary Table 6 | List of buffer compositions.** The buffers were used for protein purification, reconstitution into liposomes, activity measurements, and determination of protein concentration (see *Methods*).

| **Buffer** | **Component** | **Concentration** |
|---|---|---|
| A-buffer | MES/NaOH | 50 mM, pH 6.0 |
| | NaCl | 50 mM |
| A*-buffer | MES/NaOH | 50 mM, pH 6.0 |
| | NaCl | 50 mM |
| | $MgCl_2$ | 5 mM |
| A* pH 6.8-buffer | MES/NaOH | 50 mM, pH 6.8 |
| | NaCl | 50 mM |
| | $MgCl_2$ | 5 mM |
| | Glycerol | 10% (v/v) |
| A* LMNG-buffer | MES/NaOH | 50 mM, pH 6.0 |
| | NaCl | 50 mM |
| | $MgCl_2$ | 5 mM |
| | Glycerol | 10% (v/v) |
| | LMNG | 0.005% (w/v) |
| Binding buffer | MES/NaOH | 50 mM, pH 6.8 |
| | NaCl | 50 mM |
| | $MgCl_2$ | 5 mM |
| | Glycerol | 10% (v/v) |
| | LMNG | 0.005% (w/v) |
| Elution buffer | MES/NaOH | 50 mM, pH 6.8 |
| | NaCl | 50 mM |
| | $MgCl_2$ | 5 mM |
| | Glycerol | 10% (v/v) |
| | LMNG | 0.005% (w/v) |
| | Imidazole | 20 mM |
| Liposome buffer | MES/NaOH | 50 mM, pH 6.7 |
| | NaCl | 50 mM |
| Reconstitution buffer | HEPES | 20 mM, pH 7.5 |
| | KCl | 200 mM |
| | Sucrose | 73 mM |
| | $MgSO_4$ | 5 mM |
| | L-α-phosphatidylcholine | 0.5% (w/v) |
| | n-Octylglucoside | 1.1% |
| | Sodium desoxycholate | 0.6% (w/v) |
| | Sodium cholate | 0.6% (w/v) |
| ACMA buffer | $KH_2PO_4$/KOH | 5 mM |
| | KCl | 50 mM |
| | $MgCl_2$ | 1 mM |
| Biuret reagent | NaOH | 0.2% (w/v) |
| | Potassium sodium tartrate | 0.5% (w/v) |
| | $CuSO_4$ | 0.3% (w/v) |
| | KI | 0.5% (w/v) |
| Oxonol buffer | MES/KOH | 200 mM |
| | $MgSO_4$ | 1 mM |
| | Monensin | 0.1 μM |
| | d-Mannitol | 300 mM |
| Cryo-EM buffer | MES/NaOH | 50 mM, pH 6.0 |
| | NaCl | 250 mM |
| | LMNG | 0.005% (w/v) |

**Supplementary Table 7 | Designed primers and plasmids.**

| **Primers / Plasmid** | **Sequence** |
|---|---|
| NuoH Y156F$^H$ fwd | gcgcagaccctAagcTTCgaagtgttcctcggg |
| NuoH Y156F$^H$ rev | cccgaggaacacttcGAAgctTagggtctgcgc |
| NuoH G301L$^H$ fwd | gtccttcCTctggaaaatctgcctgccgctgacgctgatcaac |
| NuoH G301L$^H$ rev | gattttccagAGgaaggacattacctggtcataacgcggacgcggtaacg |
| NuoH H208A$^H$ fwd | ggtatgtGCccgtcacccgtttgaccag |
| NuoH H208A$^H$ rev | cgggtgacggGCacataccgccacgcccgcgatg |
| pBAD*nuo*His*nuoF* | cam$^R$, orip$_{15a}$, *araC*, P$_{araBAD}$, *nuoA-N*, $_{His}$*nuoF* |

**Supplementary Table 8 | NADH:DQ oxidoreductase activity of *E. coli* Complex I (WT) and variants.** Specific activities are reported as mean ± standard deviation of triplicate measurements of two biological replicates. Activities are additionally expressed as percentages relative to the parent Complex I (WT), which was set to 100%.

| **Strain** | **NADH:DQ oxidase activity** | |
|---|---|---|
| | **(U mg$^{-1}$)** | **(%)** |
| WT | 2.695 ± 0.05 | 100 ± 2.2 |
| Y156F$^{H}$ | 2.255 ± 0.09 | 83.6 ± 3.2 |
| G301L$^{H}$ | 1.657 ± 0.23 | 60.3 ± 8.9 |
| Y156F$^{H}$/ G301L$^{H}$ | 0.665 ± 0.13 | 24.7 ± 6.1 |
| H208A$^{H}$ | 0.658 ± 0.17 | 24.3 ± 6.0 |

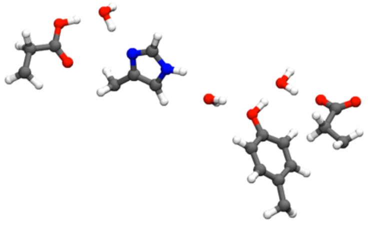

**Supplementary Movie 1 | Proton transfer along the E-channel and the Tyr156$^{H}$ flip.**

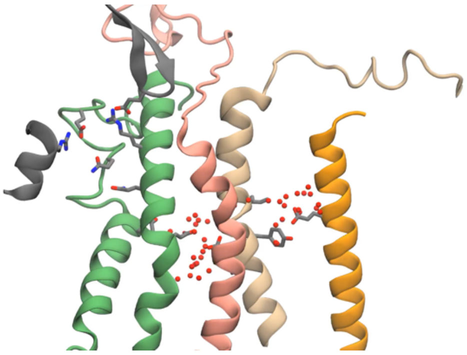

**Supplementary Movie 2 | MD simulations of the conformational flip of Tyr156$^{H}$.**
